# POLYGALACTURONASES REGULATED BY AUXIN facilitate root cell elongation in *Arabidopsis thaliana* via pectin remodeling

**DOI:** 10.1101/2025.05.07.652666

**Authors:** Monika Kubalová, Anna Kampová, Stanislav Vosolsobě, Karel Raabe, Brigita Simonaviciene, Yoselin Benitez-Alfonso, Karel Müller, Eva Medvecká, Matyáš Fendrych

**Affiliations:** Institute of Experimental Botany of the Czech Academy of Sciences, 16502 Prague, Czech Republic; Department of Experimental Plant Biology, Charles University, 12844 Prague, Czech Republic; Centre for Plant Sciences. Faculty of Biological Sciences, University of Leeds, Leeds, United Kingdom

## Abstract

Root cell elongation, the main driver of root growth, is tightly associated with cell wall remodeling, particularly through pectin modifications, which facilitate cell wall loosening and strengthening while maintaining structural integrity. Root cell elongation is precisely regulated by the phytohormone auxin, which has long been known to inhibit this process. The molecular pathways through which auxin influences cell wall modifications remain poorly understood.

In this study, we explore the transcriptional regulation of cell wall-related genes by auxin in *Arabidopsis thaliana* roots. The nuclear auxin pathway altered the expression of numerous cell-wall related genes, suggesting dynamic modification of the cell wall during root cell elongation. We identified novel root-specific polygalacturonases (PGs), enzymes involved in pectin degradation, which we termed *POLYGALACTURONASES REGULATED BY AUXIN (PGRAs). PGRAs* are expressed specifically in the root epidermis, beginning at the elongation zone.

Our results demonstrate that induction of *PGRA1* expression initially promotes root cell elongation, while long term overexpression inhibits root growth. Auxin downregulates *PGRA1* in the elongation zone, and plants lacking *PGRAs* fail to increase root growth rate in response to reduced auxin levels. This suggests that auxin downregulates *PGRA* expression to prevent PGRA-mediated pectin remodeling, thereby contributing to inhibition of root cell elongation. We established a novel link between auxin signaling and pectin modifications in the control of cell growth.

These findings provide new insights into the molecular mechanisms through which auxin regulates root cell elongation, highlighting the role of pectin matrix modifications in this process.

## Introduction

Root cell elongation drives root growth, allowing plants to explore soil for nutrients and water while adapting to environmental changes. This essential process is tightly regulated by the phytohormone auxin (Overvoorde et al., 2010; Roychoudhry & Kepinski, 2022). While not fully understood, nuclear, cytoplasmic, and apoplastic auxin signaling pathways all contribute to the regulation of cell elongation (Dubey et al., 2023; Kubalová et al., 2024, 2025; L. Li et al., 2021; Serre et al., 2021, 2023). Nuclear auxin pathway (NAP) relies on transcriptional changes in the nucleus - auxin is perceived by the TRANSPORT INHIBITOR RESPONSE1/AUXIN-SIGNALLING F-BOX (TIR1/AFB)-AUXIN/INDOLE-3-ACETIC ACID (Aux/IAA) receptor complex, leading to the degradation of Aux/IAA transcriptional repressor proteins. This releases AUXIN RESPONSE FACTORS (ARFs) to activate or repress target genes (Leyser, 2018).

One of the key manifestations of auxin’s role in regulating root cell elongation is the gravitropic response. Upon gravistimulation, auxin accumulates in the lower part of gravistimulated root, inhibiting cell elongation via a combination of transcription-dependent and non-transcriptional processes (Dubey et al., 2023; Kubalová et al., 2024; L. Li et al., 2021; Serre et al., 2021, 2023). The reduced auxin levels in the upper part of gravistimulated root allow cell elongation, causing the root to bend in the direction of gravity (Friml et al., 2002).

Cell growth depends on the balance between internal turgor pressure (Dünser et al., 2019) and the strength of the cell wall (CW). The primary CW consists of polysaccharides cellulose, hemicellulose (with the most common xyloglucan in angiosperms), pectins and proteins, such as arabinogalactan proteins, expansins (EXPs) and extensins (EXTs), and other compounds (Fry, 1988). The plant CW protects the cell, provides mechanical support, connects cells into tissues and organs, and regulates intercellular communication. To fulfill these roles, the CW must continuously balance the need for structural support with the ability to expand (Daniel J. Cosgrove, 2022).

The extensibility of the CW determines the extent of cellular expansion (Lintilhac, 2014). During cell elongation, the CW is loosened by breaking down and reassembling CW components, allowing the cell to expand (D. J. Cosgrove, 1993; Daniel J. Cosgrove, 2022).

It has been known for decades that auxin application modulates CW properties (Nishitani & Masuda, 1981), however, the exact molecular mechanisms linking CW changes and auxin-modulated root growth are not fully understood yet. Auxin regulates the biophysical properties of the CW by modulating the expression and activity of enzymes that are involved in CW biosynthesis and remodeling (Barbez et al., 2017; Jonsson et al., 2021; Majda & Robert, 2018). The status of pectin is vital in this process (Barbez et al., 2017; Jobert et al., 2023).

Pectins, a group of complex polysaccharides, play an essential role in the structural integrity of plant CWs and contribute to their physical properties, such as porosity and elasticity by forming hydrogels and interacting with other CW components (Daniel J. Cosgrove, 2022; Voragen et al., 2009). The most abundant pectin in the primary CW of *Arabidopsis thaliana* is homogalacturonan (HG) - a linear polymer of α-(1,4)-D galacturonic acid (Zablackis et al., 1995). HGs are modified by various enzymes for balancing CW stiffness and plasticity, supporting growth, facilitating stress responses, and maintaining structural and functional integrity of plant tissues (Chebli & Geitmann, 2017; Duan et al., 2020; Haas et al., 2020; Molina et al., 2021; Wachsman et al., 2020; Zamil & Geitmann, 2017). Important modification of pectin matrix is methylesterification (Wormit & Usadel, 2018). Once highly methylesterified pectin is transported to the CW, the HG undergoes selective enzymatic demethylation by wall-bound pectin methylesterases (PMEs), the activity of which is regulated by pectin methylesterase inhibitors (PMEIs). PME activity results in the formation of free carboxyl groups (Wormit & Usadel, 2018). According to the egg-box model, demethylated adjacent HG are bridged by calcium ions, leading to a stiffer and less extensible wall (Cao et al., 2020; Vincken et al., 2003). Additionally, low methylesterification of HG generates cleavage sites for pectin-degrading enzymes - pectinases, resulting in HG depolymerization and the loosening of the CW (Hocq et al., 2020; Xiao et al., 2014).

One class of pectin-degrading enzymes are polygalacturonases (PGs), which hydrolyze HG backbones. PGs can be classified into two groups based on their mode of action, with exo-PGs cleaving at the chain ends and endo-PGs hydrolyzing internal glycosidic bonds. Arabidopsis PG gene family consists of 68 genes (González-Carranza et al., 2007; Kim et al., 2006) which play a roles in various cellular processes, such as cell proliferation, expansion, morphogenesis, and separation (Hocq et al., 2020; Leng et al., 2017; Ogawa et al., 2009; Rhee et al., 2003; Rui et al., 2017; Sun et al., 2020; Vogel et al., 2002; Xiao et al., 2017, 2014; Yang et al., 2021). Interestingly, a recent study proposed that the interplay between PME and PG activities balances cell expansion and separation (Barnes et al., 2022).

Auxin regulates the expression and activity of many pectin modifying enzymes (Jobert et al., 2021; Kubalová et al., 2024; Lewis et al., 2013; Nemhauser et al., 2006). Although it is clear that the dynamic interaction between auxin and pectin is critical for plant development and physiology (Jobert et al., 2023), molecular mechanisms and biological significance of their interactions remain poorly understood.

We aimed to reveal the mechanism by which the NAP regulates Arabidopsis root cell elongation, focusing on CW modifications. We combined transcriptome profiling of root cells with biochemical analysis of CW and imaging techniques for cellular resolution of pectin matrix and characterized a novel group of pectin-degrading enzymes – PGs. Using a genetic approach, we described PGs’ role in NAP-regulated root cell elongation. We showed that NAP negatively regulates the expression of PGs and that their expression is essential to allow roots to grow under suppressed auxin conditions.

## Results

### 1. Nuclear auxin pathway induces transcriptional changes of cell wall-related genes during modulation of root cell elongation

To understand transcriptional changes governed by the nuclear auxin pathway (NAP) during root cell elongation, we re-analyzed our time-resolved transcriptomic profiling dataset (Kubalova et al., 2025) of root cells overexpressing dominant versions of auxin transcriptional repressor *Aux/IAA17 - AXR3-1* or dominant version of auxin transcriptional activator ARF5/MP - *ΔMP*. We compared transcriptomic profiles of cells undergoing enhanced elongation due to repression of the auxin transcriptional pathway (pG1090::XVE>>*AXR3-1*) with those of cells exhibiting inhibited root cell elongation due to activated auxin signaling (pG1090::XVE>>*ΔMP*) (Fig.1A,B; Fig.S1A,B).

**Figure 1:**
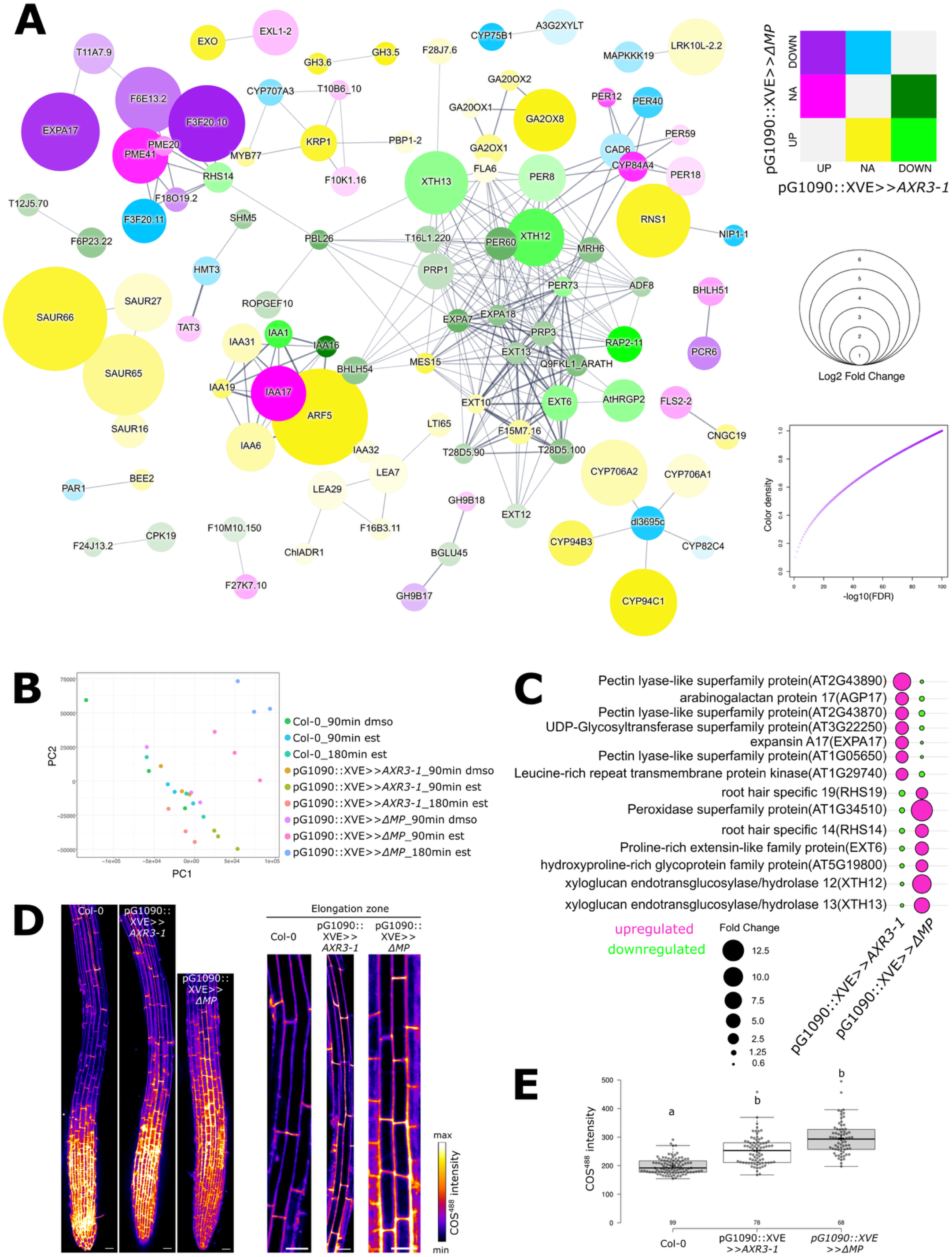
*AXR3-1* and *ΔMP* show an opposite regulation of cell wall remodeling genes. A) STRING network of differentially expressed genes after 180-minute estradiol induction vs. 90-minute DMSO control treatment in pG1090::XVE>>*ΔMP* or pG1090::XVE>>*AXR3-1* lines (*FDR* ≤ 0.05 and |*fold change*| ≥ 2 in at least one comparison). Color coding indicates significant up- or downregulation in one or both lines. NA=no change. Circle diameter and color intensity represent *log2FC* and √-log10(*FDR*), respectively, with the higher values from either line displayed. Connection lines indicate the confidence of all interaction types except text-mining. Non-connected genes are omitted from the interaction network. UP=upregulated, DOWN=downregulated, NA=no change. B) Quality control of processed RNA-seq data visualized by Principal Component Analysis generated by sleuth R package (v 0.30.1). C) Differentially expressed genes associated with the cell wall, after 180 min of estradiol treatment compared to 90 min of DMSO treatment in pG1090::XVE>>*ΔMP* or pG1090::XVE>>*AXR3-1* lines. The circle diameter indicates the fold change, with magenta representing upregulation and green representing downregulation. D) Root tips (left) and epidermal cells in elongation zone of Col-0, pG1090::XVE>>ΔMP or pG1090::XVE>>AXR3-1 treated with estradiol for 4 h and stained with COS488 for 30 min. Scale bar = 50 μm. E) Quantification of the COS^488^ intensity measured in the longitudinal cell walls of individual epidermal cells in the elongation zone of Col-0, pG1090::XVE>>*ΔMP* or pG1090::XVE>>*AXR3-1* treated with estradiol for 4 h and stained with COS^488^ for 30 min. The number of measured individual data points is indicated. F) Different letters indicate statistically significant differences between groups based on Kruskal-Wallis-test followed by post-hoc Dunn’s test.

As expected, induction of auxin signalling by accumulation of its transcriptional activator *ΔMP* led to activation of auxin early inducible genes, such as Aux/IAA proteins, SAURs, GH3 (Bao et al., 2024) or GA2OXs (Kubalova et al., 2025). The pronounced alterations in CW-related genes indicate modifications in CW dynamics (Fig.S1A,B).

Upon *AXR3-1* induction, several genes encoding *PGs, PMEs* and *PMEIs* exhibited increased expression, while other members of these groups showed decreased expression. Similarly, *AXR3-1* induction led to both upregulation, resp. downregulation of different members belonging to the *EXP* gene family. In contrast, genes involved in the xyloglucan metabolic process and those regulating *EXTs* were predominantly downregulated (Fig.S1B,C). Conversely, induction of *ΔMP* led to upregulation of genes associated with the xyloglucan metabolic process and those encoding *EXTs*, whereas genes encoding PG enzymes were downregulated. Members of the *EXP* and *PME* or *PMEI* gene families displayed both upregulation and downregulation (Fig.S2A,B). These findings suggest that auxin-regulated root cell elongation is controlled through the modulation of xyloglucan, EXTs, EXPs, and the pectin matrix.

We focused on genes exhibiting opposing expression patterns between rapidly elongating pG1090::XVE>>*AXR3-1* roots and inhibited pG1090::XVE>>*ΔMP* roots (Fig.1A,C; Fig.S3A), because their expression level might correlate with the cell elongation phenotype. The most prominent genes were involved in the xyloglucan metabolic process and genes related to pectin metabolism. We focused on the pectin matrix as a key component of the CW affected by auxin signaling (Jobert et al., 2023).To investigate transcriptome-predicted changes, we first performed glycome profiling on whole root extracts. The results did not reveal a clear correlation between alterations in extractable pectin levels and the modulation of auxin signaling (Fig.S3B). However, tissue homogenization results in the loss of spatial information at the cellular level, potentially masking subtle, cell-type-specific changes in CW composition. Therefore, we conducted a more detailed analysis of pectins in the root elongation zone, where active cell expansion occurs and CW modifications are expected to be most pronounced. To stain demethylesterified pectins in living roots, we used polysaccharide-based probe - positively charged and fluorescently tagged chitosan oligosaccharide (COS^488^) probe (Mravec et al., 2014). Compared to the control, repression of auxin signaling led to a mild increase in the demethylesterified pectins, whereas activation of auxin signaling resulted in a more significant increase in their abundance in root elongation zone (Fig.1D,E). Changes in pectin levels can result from altered degrees of methylesterification, as well as from the regulation of pectin biosynthesis or degradation.

Further, we focus on pectin-modifying enzymes - polygalacturonases (PGs). Focusing on these genes is particularly interesting because PGs directly catalyze pectin degradation in the primary CW, a key step in CW loosening important for cell elongation (Rui et al., 2017; Verhage, 2021; Xiao et al., 2014).

### 2. PGRAs are pectin degrading enzymes that are negatively regulated by nuclear auxin pathway

Among the set of pectin-modifying enzymes, a yet-uncharacterised group of 3 PGs displayed the most striking expression pattern (Fig.2A). We named these genes *POLYGALACTURONASE REGULATED BY AUXIN* (*PGRA*). The expression of all 3 *PGRA* genes was downregulated upon induction of pG1090::XVE>>*ΔMP* and upregulated in roots with induced pG1090::XVE>>*AXR3-1*. This led us to hypothesize that auxin negatively regulates *PGRAs* expression, thereby preventing the CW loosening required for root cell elongation. For further investigation, we primarily focused on *PGRA1*, as it exhibited the highest responsiveness to auxin signalling modulation while also being highly expressed in the root (Fig.2A,B).

**Figure 2:**
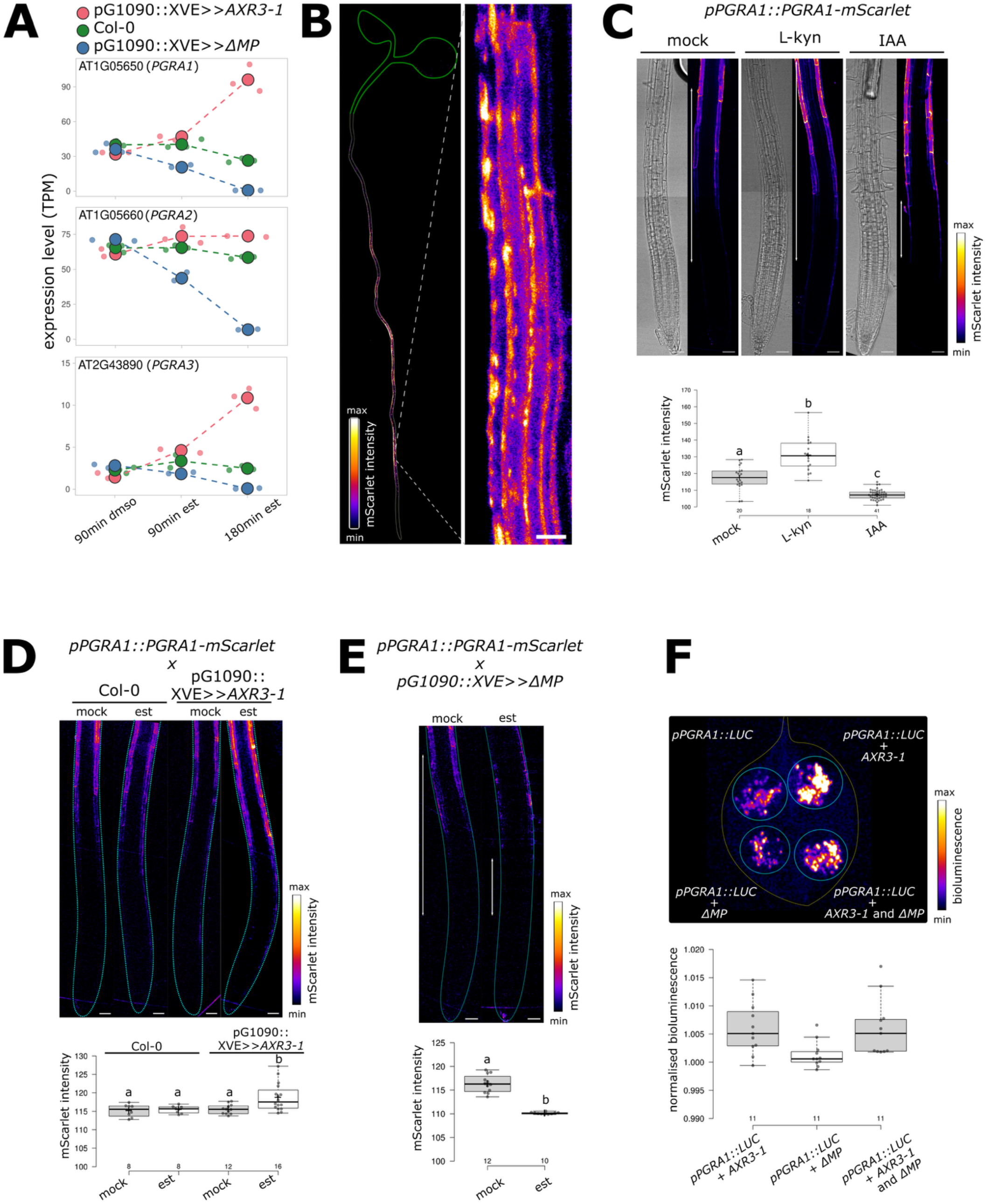
*PGRA1* gene is expressed in root and is negatively regulated by auxin nuclear pathway. A) Relative expression level of *PGRA1, PGRA2* and *PGRA3* represented as TPM in pG1090::XVE>>*AXR3-1*, pG1090::XVE>>*ΔMP* and Col-0 root cells after 90 min of DMSO or 90 or 180 min of estradiol treatment. B) Expression of *pPGRA1::PGRA1-mScarlet* in Arabidopsis seedlings (left), with a focus on root epidermal cells (right). Scale bar = 10μm. C) Root tip of *pPGRA1::PGRA1-mScarlet* treated with 1.5 μM L-kyn for 4h or 50 nM IAA for 6h. The boxplot shows the mScarlet intensity measured in the longitudinal cell walls of individual epidermal cells in the elongation zone. The number of measured individual data points is indicated. White arrows indicate the elongation zone. Scale bar = 50μm. D) Root tip of *pPGRA1::PGRA1-mScarlet* crossed with pG1090::XVE>>*AXR3-1* or Col-0 and treated with estradiol for 4h. The boxplot shows the mScarlet intensity measured in the longitudinal cell walls of individual epidermal cells in the elongation zone. Turquoise line marks the outline of the root. Scale bar = 50μm. E) Root tip of *pPGRA1::PGRA1-mScarlet* crossed with pG1090::XVE>>*ΔMP* and treated with estradiol for 16h. The boxplot shows the mScarlet intensity measured in the longitudinal cell walls of individual epidermal cells in the elongation zone. White arrows indicate the elongation zone, and the turquoise line marks the outline of the root. Scale bar = 50μm. F) Activity of *pPGRA1::LUC* in tobacco leaf co-infiltrated with/without p35S::*ΔMP-mScarlet* and p35S::*AXR3-1-mVenus*. The boxplot shows the luminescence of *pPGRA1::LUC* co-infiltrated with/without p35S::*ΔMP-mScarlet* and p35S::*AXR3-1-mVenus*. Normalised to luminescence of *pPGRA1::LUC* without co-infiltration. Different letters indicate statistically significant differences between groups based on one-way ANOVA followed by Tukey HSD (C,D) or Student t-test (E).

To confirm the transcriptomic data, we generated a reporter line in which the *PGRA1* tagged with mScarlet was driven by its native promoter (*pPGRA1::PGRA1-mScarlet*). PGRA1-mScarlet signal was detected specifically in the root. The PGRA reporter showed a specific expression pattern in the root epidermis, with increased expression starting in the elongation zone (Fig.2B). This line was treated with the auxin Indole-3-acetic acid (IAA) or L-kynurenine (L-Kyn), an auxin biosynthesis inhibitor (He et al., 2011). Additionally, we crossed this reporter line with pG1090::XVE>>*AXR3-1* and pG1090::XVE>>*ΔMP* lines. Inhibition of auxin biosynthesis, as well as induction of *AXR3-1* expression, significantly increased PGRA1-mScarlet accumulation in the elongation zone (Fig.2C,D). In contrast, activation of auxin signaling by *ΔMP* induction or IAA treatment decreased PGRA1-mScarlet accumulation in the elongation zone (Fig.2C,D). Interestingly, the PGRA1-mScarlet accumulation in the differentiation zone was not affected by these treatments. These results demonstrate that *PGRA1* expression is negatively regulated by NAP in the epidermal cells of the root elongation zone. Additionally, we confirmed that *AXR3-1* and *ΔMP* regulate the *PGRA1* promoter activity using the heterologous *Nicotiana benthamiana* expression system. *LUCIFERASE* gene (*LUC*) driven by *PGRA1* promoter was co-expressed with the *AXR3-1* tagged with mVenus and/or with *ΔMP* tagged with mScarlet (Fig.2F). The expression of the constructs was verified using confocal microscopy (Fig.S4). The co-expression with *AXR3-1* significantly increased the activity of *PGRA1* promoter while *ΔMP* had no effect (Fig.2F). Interestingly, *PGRA* promoters do not contain binding sites for activating ARFs; only the *PGRA1* promoter contains a binding site for ARF1 (tgtctc), which is considered an ARF with a repressive effect on transcription (Tiwari et al., 2003).

To characterize the biochemical functions of *PGRAs*, we first utilized bioinformatic tools. Structural comparison of PGRA1 (which is 89.1% identical to PGRA2) with previously characterized plant PGs revealed that PGRAs contain a glycosyl hydrolase family 48 (Glyco48) domain, typical for PG enzymes (Fig.3A). Phylogenetic analysis (Fig.S5A,B) or structural prediction analysis (Fig.S5C,D) does not allow for conclusive characterization of these enzymes as either exo-PG or endo-PG.

**Figure 3:**
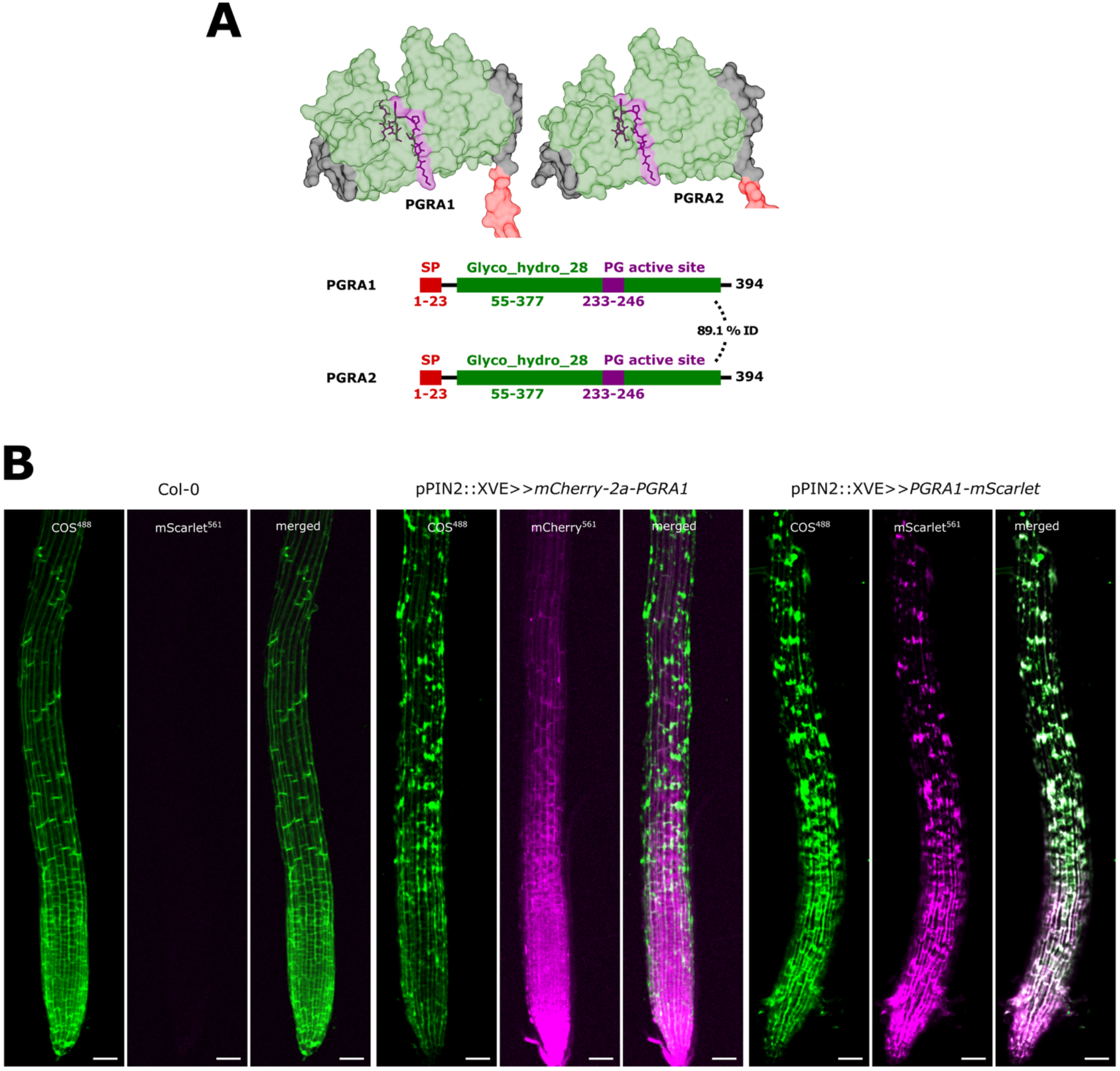
*PGRA1* is a pectin degrading enzyme. A) Surface structures of PGRA1 and PGRA2 protein models (upper part) and their domain organization and protein sequence identity (lower part). Surface structure coloring follows the schematic model: SP=signal peptide (red), GH28 domain (green), and conserved PG active site (purple) with visible atom structure. B) Root tips of Col-0, pPIN2::XVE>>*mCherry-2a-PGRA1* and pPIN2::XVE>>*PGRA1-mScarlet* grown on estradiol for 5d and stained with COS^488^ for 30 min. Scale bar = 50 μm.

To validate this prediction, COS probe was used to stain demethylated pectins in plants overexpressing *PGRA1*. We used an estradiol-inducible system, where *PGRA1* gene is expressed in the outer root tissues using the PIN2 promoter. To eliminate the potential impact of PGRA protein tagging on its function, we analyzed both C-tagged (pPIN2::XVE>>*PGRA1-mScarlet*) and untagged (pPIN2::XVE>>*mCherry-2a-PGRA1*) *PGRA1* variants, which yielded identical results. Compared to control, COS staining revealed altered pectin distribution, showing delocalization of the demethylated pectins in *PGRA1*-overexpressing roots (Fig.3B). Interestingly, PGRA1 colocalized with the sites of pectin accumulation. Taken together, these data demonstrate that *PGRAs* encode functional PG enzymes that are negatively regulated by NAP.

### 3. PGRAs promote root cell elongation

To examine the biological role of *PGRAs* in root development, we analyzed the root growth rate in plants with increased *PGRA* expression. Using an estradiol-inducible system, we monitored root growth dynamics following *PGRA1* induction (Fig.4A). Induction of both tagged (pPIN2::XVE>>*PGRA1-mScarlet)* or untagged (pPIN2::XVE>>*mCherry-2a-PGRA1*) *PGRA1* expression initially promoted root elongation, indicating a positive effect on root cell expansion. However, prolonged *PGRA1* overexpression led to root growth inhibition (Fig.4B). Consistent with this, plants with constitutive *PGRA1* overexpression (pG1090::*PGRA1-mScarlet)* displayed a dose-dependent response: weak overexpression lines showed an extended root elongation zone, whereas strong overexpression caused its inhibition (Fig.4C). Negative effect of overexpressed *PGRA1* is likely due to an activated stress response triggered by excessive *PGRA1* activity as suggested by transcriptomic analyses. Accumulation of PGRA1 protein induced the expression of stress and defense related genes (Fig.S6A,B, S7A). These findings suggest that the primary function of PGRA is to hydrolyze demethylesterified pectin in the CWs, leading to promotion of root growth.

**Figure 4:**
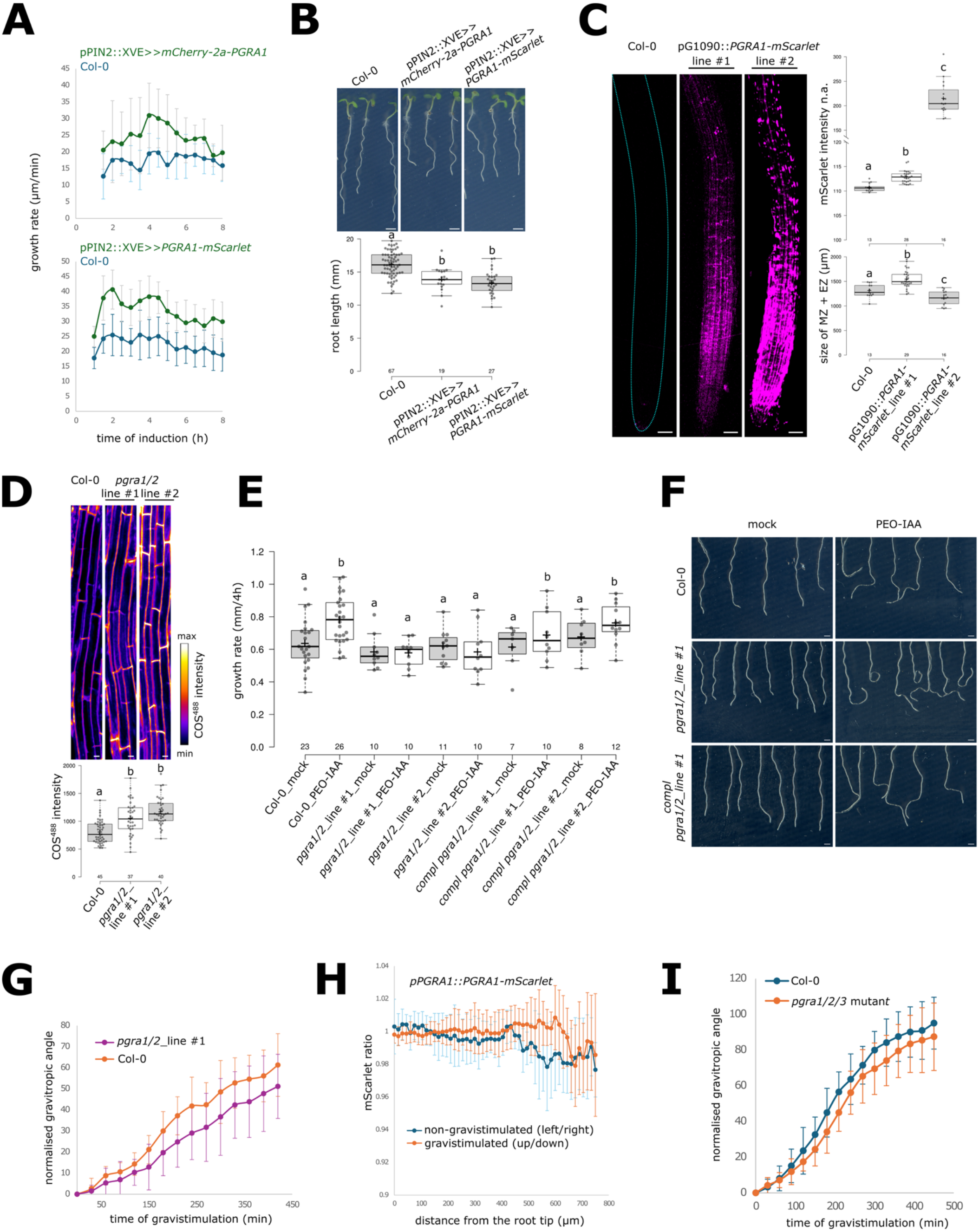
*PGRA*1 promotes root cell elongation and is essential for growth promotion under low auxin conditions. A) Growth rate of pPIN2::XVE>>*mCherry-2a-PGRA1* and pPIN2::XVE>>*PGRA1-mScarlet* after estradiol treatment, corresponding to time of induction. n≥ 8. B) Col-0, pPIN2::XVE>>*mCherry-2a-PGRA1* and pPIN2::XVE>>*PGRA1-mScarlet* grown on estradiol for 5d. The boxplot shows the quantification of root length. The number of measured individual data points is indicated. Scale bar = 0.1 cm. C) Root tips of pG1090::*PGRA1-mScarlet*. The boxplot shows the mScarlet intensity measured in the elongation zone and the distance from the root tip to the end of elongation zone. (EZ=elongation zone, MZ=meristematic zone). Turquoise line marks the outline of the root. Scale bar = 50 μm. D) Epidermal cells of *pgra1/2* mutants stained with COS^488^. The boxplot shows the COS^488^ intensity measured in the longitudinal cell walls of individual epidermal cells in the elongation zone. The number of measured individual data points is indicated. Scale bar = 10 μm. E) Growth rate of *pgra1/2* mutants and its complementation line treated for 4 h with 10 μM PEO-IAA. F) Root tips of *pgra1/2* mutants and its complementation line treated for 24 h with 10 μM PEO-IAA. Scale bar = 0.1 cm. G) Normalized gravitropic angle of *pgra1/2* mutant. n≥ 10. H) mScarlet excitation ratio of left/right epidermal layer in non-gravistimulted root or mScarlet excitation ratio of up/down epidermal layer in gravistimulated root of *pPGRA1::PGRA1-mScarlet*. n≥ 9. I) Normalized gravitropic angle of *pgra1/2*/3 mutant. n≥ 10. Different letters indicate statistically significant differences between groups based on one-way ANOVA followed by Tukey HSD.

Next, we generated 2 independent alleles of *pgra1/2 double* mutants using CRISPR/Cas9, and crossed it with *pgra3* T-DNA insertion line to create *pgra1/2/3 triple* mutant (Fig.S8A). Mutation in these genes led to increased level of demethylated pectins in longitudinal CW of cells in the elongation zone (Fig.4D; Fig.S8B). Based on prior findings, we hypothesized that under low auxin conditions, *PGRAs* are upregulated to promote CW loosening and enhance root cell elongation.

To test this, we treated *pgra1/2 double* mutant with PEO-IAA to inhibit auxin signaling. While control plants showed increased root cell elongation under this condition, *pgra1/2 double* mutants did not (Fig.4E). Notably, growth rate of *pgra1/2 double mutant* under mock conditions is comparable to control, indicating that *PGRAs* are essential for PEO-IAA-induced root growth promotion. Interestingly, after long-term PEO-IAA treatment, *pgra1/2 double* mutants displayed a more agravitropic phenotype compared to control plants (Fig.4F).

To evaluate the relevance of the auxin-PGRA interaction in a physiological context, we analyzed the gravitropic response of *pgra1/2 double* mutants. Gravitropic response of *pgra1/2 double* was slightly slower than gravitropic response of control (Fig.4G). However, we were unable to observe a differential expression of *PGRA1* between the upper and lower sides of gravistimulated roots (Fig.4H). To overcome potential gene redundancy, we prepared *pgra1/2/3* triple mutant. Interestingly, gravitropic response of *pgra1/2/3* triple mutant was comparable to control plants (Fig.4I). Similar to *pgra1/2 double* mutant, *pgra1/2/3 triple* mutant showed resistance to PEO-IAA treatment (Fig.S8B).

Since sugar metabolism strongly affects CW properties and the presence of saccharose in the medium could prevent the expression of weaker phenotypes in CW mutants (Dash et al., 2023; Y. Li et al., 2007; Pottier et al., 2023), we tested *pgra* mutants grown and treated without adding saccharose to the medium. However, their gravitropic response under these conditions remained the same - *pgra1/2 double* was slightly slower, while the response of *pgra1/2/3* triple mutant was comparable to control (Fig.S8C).

Taken together, these findings indicate that PGRA-induced remodeling of pectin matrix plays a critical role in promoting root cell elongation and *PGRA* genes are negatively regulated by the NAP in the root elongation zone.

## Discussion

CW modifications, especially through the remodeling of pectin, play a critical role in facilitating the expansion of cells. Despite the well-established role of auxin in controlling root growth, how auxin regulates the expression of enzymes involved in CW remodeling during root cell elongation is not understood.

In this study, we addressed this knowledge gap by investigating the transcriptional regulation of CW-related genes by auxin, focusing specifically on pectin modifications during root cell elongation. We characterised novel PGs - PGRA enzymes, which are expressed in root and are negatively regulated by the NAP in the elongation zone of the root. These enzymes are essential for promoting root cell elongation under conditions where auxin signalling is suppressed.

PGs fine-tune cell elongation, expansion and adhesion through modulation of pectin matrix (Safran et al., 2023; Xiao et al., 2014) In Arabidopsis roots, PGs effect is primarily associated with promoting growth (Hocq et al., 2020; Nagayama et al., 2022; Rui et al., 2017; Xiao et al., 2014), but some studies showed negative effect on it (Safran et al., 2023; Xiao et al., 2017). These differences may be attributed to the extent of deregulated PG activity, as supported by our data. Overexpression of here described *PGRAs* initially leads to the promotion of root cell elongation, likely caused by the loosening of the CW through pectin cleavage and its relocalization. PGRA enzymes are specifically expressed in the root and at the cellular level, they are localized into specific domains. The localized modification of pectin in microdomains is thought to be more effective in coordinating specific responses to external signals (Dauphin et al., 2022). Interestingly, long-term overexpression of PGRA also resulted in its localization within specific domains. Since the accumulation domains of PGRA overlap with demethylated pectin accumulation, it can be inferred that the PGRA enzyme accumulates at the site of its substrate.

To investigate the molecular function of PGRAs, we utilized homology modeling and compared the predicted structures to the experimentally validated structures of known endo/exo-PGs; however, we were unable to determine whether PGRAs function as exo- or endo-PGs. Due to structural differences, exo- and endo-PGs are often proposed to have distinct roles in the CW (Henrissat & Davies, 1997; Zhang et al., 2007), yet their specific functions in growing cells remain poorly understood. Since exo-PGs cleave only one to two residues at a time (Abbott & Boraston, 2007), their impact on CW mechanics remains uncertain. In contrast, previously characterized endo-PGs have been directly linked to cell expansion (Xiao et al., 2017, 2014), suggesting that PGRAs may also function as endo-PGs.

Mutation of *PGRA* genes significantly increased the levels of demethylated pectins in the elongation zone but did not affect root growth under normal conditions. However, the lack of response in *PGRA*-deficient plants to reduced auxin signaling and their negative regulation by NAP suggests *PGRAs’* role in auxin-regulated growth. These results indicate that *PGRA* genes are important for promoting elongation when auxin levels or signaling are low. Plants lacking 2 *PGRA* genes showed a mild delay in gravitropic response, suggesting that their activation in the upper part of gravistimulated root contributes to the gravitropic bending. However, despite auxin accumulation on the lower side of the root and depletion on the upper side during gravistimulation (Friml et al., 2002), PGRA1-mScarlet signal intensity gradient could not be detected between upper and lower side of gravistimulated root. This could be due to the fact that the decrease in auxin levels is not large or sustained enough to significantly reduce PGRA1-mScarlet levels to a measurable value. Nevertheless, only slight delay in the gravitropic response suggests that *PGRA* genes do not play a significant role in differential growth during gravitropism, and their role in auxin-regulated processes in a natural context remains to be defined.

We expected that the mutation of an additional *PGRA3* gene would result in a more severe phenotype. However, the *pgra1/2/3 triple* mutant displayed a weaker phenotype compared to the *pgra1/2 double* mutant. This observation may be attributed to a compensatory mechanism, where upregulation of other PGs could alleviate the phenotypic effects of the triple mutation, suggesting partial compensation through the activation of other PG family members, as shown in previous studies (Hocq et al., 2020; Kim & Patterson, 2006). Testing this hypothesis would require assessing the expression levels of additional PGs in the *pgra1/2/3 triple* mutant.

The negative regulation of *PGRAs* by auxin is particularly intriguing. *ARF5/MP* is generally considered an activator of auxin-regulated genes (Cancé et al., 2022); however, transcriptomic data indicate that *ARF5/MP* suppresses *PGRAs* expression. Although we demonstrated a positive effect of *AXR3-1* on *PGRA1* promoter activity, the direct inhibition of the *PGRA1* promoter by *ΔMP* has not been shown. This is consistent with the absence of ARF5/MP-binding site in *PGRA1* promoter and suggests an indirect effect. ARF5/MP may activate and AXR3-1 inhibit the transcription of a *PGRA1* inhibitor. However, *PGRAs* genes are downregulated by IAA treatment within just 20 minutes (Kubalová et al., 2024), a timeframe likely insufficient for the transcription of an inhibitor and its subsequent effect on *PGRA* gene expression. *PGRA1* may be negatively regulated by auxin-induced activation of ARF1, which is thought to have a both positive and negative role on auxin induced genes (Cancé et al., 2022). While *PGRA1* contains one ARF-binding site, *PGRA2* and *PGRA3* lack such sites in their promoters, implying at least partial ARF-independent regulation.

PGs activity is closely related to pectin-methylesterification status. We demonstrated that the transcription of numerous PME and PMEI is altered by the NAP during regulation of root cell elongation. The distribution of methyl groups along the HG backbone is known to be a key determinant of CW stiffness (Braybrook & Peaucelle, 2013). Modulation of PME and PMEI activity in Arabidopsis roots has been shown to have varying effects on root growth. Both increased and decreased PME activity have been associated with longer roots in different *pme* and *pmei* mutants and overexpressors (Hewezi et al., 2008; Mase et al., 2023; Sénéchal et al., 2014; Wolf et al., 2012). Additionally, decreased pectin methylesterification in certain *PMEI* overexpressors had no impact on root length (Rößling et al., 2024), suggesting that the regulation of pectin methylesterification does not have a simple linear effect on root elongation. Specific PME-PMEI interactions (Jeong et al., 2018) or PME-PGs activity balancing (Barnes et al., 2022), rather than overall PME activity levels, may play a key role in modulating root growth. Adding additional complexity, the effect of auxin-modulated methylesterification appears to be time dependent (Gallemí et al., 2022) and tissue-specific. During apical hook development, elevated auxin levels in the inner hypocotyl increase methylesterification, restricting growth (Jonsson et al., 2021). In contrast, during organ initiation, auxin stimulates demethylesterification to soften tissues (Peaucelle et al., 2011). Previously (Kubalová et al., 2024), we investigated the mutants of the most deregulated PME and PMEI genes within our datasets, and found no significant effect on root growth, possibly due to genetic redundancy.

We showed that *PMEs* and their inhibitors *PMEIs* exhibit both upregulation and downregulation in response to auxin signaling modulation. Additionally, we observed that the level of demethylesterified pectins in the root elongation zone increases under both auxin activation and repression. These findings suggest that auxin-modulated pectin methylesterification does not have a direct, linear correlation with the modulation of root cell elongation. Tissue type, developmental stage, and cellular age are probable determinants of this relationship. Based on the transcriptomic data, we can hypothesize that specific PME isoforms selectively generate either blockwise demethylesterified HG, which likely facilitates calcium bridge formation and CW stiffening, or randomly demethylesterified HG, which creates cleavage sites for pectin-degrading enzymes, leading to CW loosening (Hocq, Pelloux, et al., 2017). We can speculate that in pG1090::XVE>>*ΔMP* roots, the increased CW stiffness due to calcium bridge formation may hinder root cell elongation. Conversely, in pG1090::XVE>>*AXR3-1* cells, enhanced pectin cleavage driven by upregulated pectinases may contribute to CW loosening and promote elongation. It is worth noting that modulation of auxin signaling by *AXR3-1* and *ΔMP* significantly altered root hair development (Kubalová et al., 2024). Therefore, the observed changes in various CW-related genes may also be attributed to the presence or absence of root hairs.

Cell elongation is largely governed by the dynamic regulation of and complex interactions among all CW components (Daniel J. Cosgrove, 2018a, 2022; Delmer et al., 2024). Consistent with this, our transcriptomic analysis revealed not only modifications in the pectin matrix but also alterations in hemicellulose composition and the deregulation of structural proteins, such as EXPs and EXTs. Pectins, EXPs, xyloglucans, and EXTs are in dynamic equilibrium, regulating the flexibility and strength of the CW (Phyo et al., 2019). Pectin status is particularly important in these interactions, with de-esterified pectin enhancing these interactions and increasing wall flexibility (Haas et al., 2021). EXPs promote CW expansion by loosening the bonds between xyloglucans and cellulose (Daniel J. Cosgrove, 2018b). Pectin methylation status also balance the activity of PGs (Barnes et al., 2022) and interaction with EXTs, likely strengthening the CW and counteracting the loosening effects (Willats et al., 2001; Wolf et al., 2009). Additionally, pectic polysaccharides (HG and rhamnogalacturonans) covalently interact with each other to form a complex network of pectin within the CW (Anderson, 2019). Therefore, changes in HG caused by the cleavage by PGRAs can affect the structure and, consequently, the function of other types of pectin matrix.

Another factor influencing the CW dynamics is apoplastic pH (Hocq et al., 2024), which is coregulated by auxin (Barbez et al., 2017; Hager et al., 1971; L. Li et al., 2021; Lin et al., 2021; Rayle & Cleland, 1970; Serre et al., 2021, 2023). Specifically, AXR3-1 expression leads to apoplast acidification, while *Δ*MP expression causes apoplast alkalinization (Kubalová et al., 2024). As CW remodeling enzymes are active in an optimal pH (Duvetter et al., 2006), apoplastic pH changes triggered by auxin may influence the overall activity of CW-modifying enzymes and the resulting effect on CW properties and cell elongation. Thus, the overall effect on CW changes is the result of a combination of altered expression and activity of CW remodeling enzymes.

Additionally, we found that long-term accumulation of PGRA1 has an inhibitory effect on growth; a prolonged or high-level overexpression of PGRA1 leads to growth suppression. This suggests that PGRA1 activity has a dual effect, depending on its concentration, or that growth inhibition results from adaptive CW mechanisms, as CW cleavage changes CW architecture (Capodicasa et al., 2004; Hocq, Sénéchal, et al., 2017; Sénéchal et al., 2014; Xu et al., 2022). Supporting this idea, transcriptomic analysis shows that plants overexpressing *PGRA1* have increased expression of stress-related, defense genes or those involved in CW modulation. Therefore, it is possible that *PGRA1* overexpression triggers stress response. Alternatively, since PGs produce short oligogalacturonides that serve as signaling molecules (Ferrari, 2013), another possible explanation is that PGRAs play a role in defense response signaling. Strong *PGRA1* expression in the epidermis of the maturation zone supports this idea. The dual role of pectin modifying enzymes has been already shown (Huerta et al., 2023).

The regulation of CW dynamics during root elongation integrates various molecular interactions and feedback mechanisms. We showed that *PGRAs* are involved in pectin modifications in plants and are transcriptionally regulated by NAP, indicating their potential role in auxin-mediated regulation of cell growth. However, the natural context of this involvement remains to be elucidated. Overall, our findings provide new insights into the role of auxin in regulating CW modifications and highlight the significance of pectin remodeling in root cell elongation.

## Acknowledgements

MF and MK received support from the European Research Council (MORpH Grant No. 101125499). MK was supported by Charles University Grant Agency (Grant No. 337021), The Endowment Fund of the Faculty of Science of Charles University and EMBO Scientific Exchange Grant (No. 10300). MF, KR and KM were supported from the project TowArds Next GENeration Crops, reg. no. CZ.02.01.01/00/22_008/0004581 of the ERDF Programme Johannes Amos Comenius. BS received a studentship from the School of Biology, Faculty of Biological Sciences, University of Leeds. YBA lab is supported by the United Kingdom Research and Innovation (UKRI) council Future Leaders Fellowship (MR/T04263X/1). The authors are grateful to Matouš Glanc for critical reading of the manuscript and Jozef Mravec for providing us with the COS probe.

## Author contribution

MF, MK - Conceptualization, Writing – Original Draft, Writing – Review & Editing, Interpretation of Results, Funding Acquisition. MK – Investigation, Validation, Visualization, Acquisition of Data, Analysis and Interpretation. MF - Supervision. SV - Formal Analysis (STRING networks), Visualization. AK, EM - Acquisition of Data, Technical Support. KR - Formal Analysis (bioinformatic analysis), Visualization. KM - Formal Analysis (transcriptomic data). BS, YBA - ELISA profiling experiments – Method Writing, Acquisition of Data, Analysis and Interpretation.

## Material and methods

### Growth conditions

Seeds were surface-sterilized by chlorine gas(10.3791/56587). After this, seeds were stratified for 2 d at 4°C. Seedlings were grown vertically on plates containing 1% (w/v) agar (Duchefa) with ½ Murashige and Skoog (MS, Duchefa, 0,5g/l MES, 1 % (w/v) sucrose, pH 5.8 adjusted with 1M KOH). Plants grown in a growth chamber with following conditions: 60% humidity, 22°C by day (16 h), 18°C by night (8 h), light intensity of 120 μmol photons m^-2 s-1^.

### Plant material and generation of transgenic plants

All lines used are in *Arabidopsis thaliana* ecotype Columbia-0 (Col-0) background.

*pgra1/2 double* mutants (AT1G05650, AT1G05660) were prepared by CRISPR/Cas9 strategy (Xing et al., 2014). Cas9 was driven by a Ubiquitin promoter and cloned into the pDGB3omega1 binary vector. gRNAs are listed in Table S1. Subsequent generations were then backcrossed with Col-0 and genotyped for homozygotes. *pgra1/2 double* mutant (version 1) was crossed with *pgra3* (GK-343D11, AT2G43890, obtained from NASC) to create *pgra1/2/3 triple* mutant. Complementant lines were prepared by crossing *pgra1/2 double* mutants with *pPGRA1::PGRA1-mScarlet* (prepared as described below). Primers used for genotyping are in Table S2. *pPGRA1::PGRA1-mScarlet* was crossed with pG1090::XVE>>*AXR3-1* (Mähönen et al., 2014) or pG1090::XVE>>*ΔMP* (Gonzalez et al., 2021).

GoldenBraid methodology (Alejandro Sarrion-Perdigones et al., 2011) was used as a cloning strategy. For stable transcriptional or translational fusion of *PGRA1*, we cloned 1739bp upstream of *PGRA1* gene and fused it with *luciferase* gene (Gould & Subramani, 1988) (*pPGRA1::LUC*) or *PGRA1* gene (including introns) tagged with mScarlet-I (*pPGRA1::PGRA1-mScarlet*). To prepare stable overexpressor lines, *PGRA1* gene (including introns) tagged with mScarlet-I was driven by G1090 promoter (Zuo et al., 2000) (pG1090::*PGRA1-mScarlet*). All these constructs were terminated by 35S terminator and cloned into alpha1 vector.

For estradiol-inducible lines, the XVE was cloned under the control of the PIN2 promoter (1.4 kb upstream of AT5G57090) and terminated by the RuBisCo terminator from *Pisum sativum*, then inserted into the alpha 1-1 vector. The *PGRA1* gene was cloned downstream of the 4xLexA operon, driven by the CaMV 35S minimal promoter (A. Sarrion-Perdigones et al., 2013) and terminated by the 35S terminator. mCherry (Shaner et al., 2004) and 2A self cleaving peptide (Samalova et al., 2006) were fused to the N-terminus of *PGRA1* (pPIN2::XVE>>*mCherry-2a-PGRA1)*, or mScarlet-I was fused to the C-terminus of *PGRA1* (pPIN2::XVE>>*PGRA1-mScarlet*), with both constructs terminated by the 35S terminator. These constructs were then inserted into the alpha1-3 vectors (Dusek et al., 2020). The transcriptional units in the alpha1-1 and 1-3 vectors were interspersed with matrix attachment regions(Dusek et al., 2020). Finally, the alpha vectors were combined with a Basta resistance cassette in the alpha2 plasmid and cloned into the pDGB3omega1 binary vector (Zuo et al., 2000). All constructs were transformed into A. thaliana ecotype Col-0 using the floral dip method (Bindels et al., 2017). Primers used for cloning are in Table S3.

### Treatments and phenotyping

To treat the plants, 5 d old plants were transferred to a treatment-containing medium. Estradiol (Sigma, 20mM stock in DMSO, 2.5 μM working concentration), Indole Acetic Acid (IAA, Sigma,10mM stock in 96% ethanol), 2-(1H-Indol-3-yl)-4-oxo-4-phenylbutyric acid (PEO-IAA, MedChemExpress, 10mM stock in DMSO) and L-kynurenine (L-kyn, Carl Roth, 3mM stock in DMSO) were used for treatments. Working concentrations and treatment time for specific experiments are given in the legend of each figure. Root growth rate was determined by measuring the distance between the positions of the root tip in successive time points. Root length was defined as the distance from the root base to the root tip. The boundary of the elongation zone was identified as the first cell that showed root hair formation, while the end of the meristematic zone was marked by the last isodiametric cell. To stain demethylated pectins, plants were treated with COS probe coupled to Alexa Fluor 488 (COS^488^) (Mravec et al., 2014) diluted 1 in 2000 in liquid ½ MS medium for 30 min, kept in dark. After washing three times, roots were transferred on ½ MS medium and imaged. Gravitropic bending was assessed by rotating plates containing seedlings by 90° and capturing images every 30 minutes. ELISA profiling of the cell wall is described in Supplemental methods.

### Microscopy, low-resolution imaging and image analysis

For high-resolution imaging, a vertical system (von Wangenheim et al., 2017) consisting of a Zeiss Axio Observer 7 microscope coupled to a Yokogawa CSU-W1-T2 spinning disk unit with 50 μM pinholes and a VSHOM1000 excitation light homogenizer (Visitron Systems) was used. Images were acquired using VisiView software (Visitron Systems, v4.4.0.14). The 561 nm laser was employed for mCherry and mScarlet (excitation 561, emission 582-636 nm), the 515 nm laser for GFP (excitation 515, emission 520-570 nm), and the 488 nm laser for COS stained samples (excitation 488, emission 500-550 nm). For low-resolution imaging, a vertically placed flatbed scanner (Perfection V700, Epson) was used. Images were acquired using Epson Scanner software (v3.9.2.1US). Image analysis was performed using ImageJ Fiji software (10.1038/nmeth.2019). The analysis of the gravistimulated roots was performed using Acorba software (Serre & Fendrych, 2022).

### Luciferase reporter assay

Luciferase reporter assay was performed as described in (Kubalova et al., 2025). *pPGRA1::LUC* was prepared as described earlier.

### Transcriptomic and gene expression analysis

Transcriptomic and gene expression analysis were done as described in (Kubalova et al., 2025). Transcriptomic data from pG1090::XVE>>*AXR3-1* and pG1090::XVE>>*ΔMP* were prepared as described in (Kubalová et al., 2024). pPIN2::XVE>>*PGRA1-mScarlet*, pPIN2::XVE>>*mCherry-2a-PGRA1* and Col-0 were grown on ½ MS plates containing 2.5 μM estradiol. 5d old roots were harvested.

The set of genes with *FDR ≤ 0.05* and |*fold change*| ≥ 2 in at least one comparison between 180-minute estradiol induction vs. 90-minute DMSO control treatment in pG1090::XVE>>*ΔMP* or pG1090::XVE>>*AXR3-1*, and in at least one of pPIN2::XVE>>*PGRA1-mScarlet* or pPIN2::XVE>>*mCherry-2a-PGRA1* line respectively, was uploaded to STRING v12.0 (https://string-db.org/) to generate an interaction diagram including all interaction types except text-mining with unconnected genes hidden. Additionally, individual diagrams for each line were generated with all genes shown and with inclusion of text-mining interaction for pG1090::XVE>>*ΔMP* and pG1090::XVE>>*AXR3-1* lines. The diagram was further edited using BASH and R to modify colors and point diameters based on transcriptomic parameters.

Gene ontology analysis was done using DAVID bioinformatics resources (Huang et al., 2009).

### Domain and structure analysis

Protein sequences for PGRA1 and PGRA2 were retrieved from UniProt under accessions AT1G05650 (UniProt ID: Q9SYK6) and AT1G05660 (UniProt ID: Q9SYK7), respectively. Domain annotations, including Glycosyl Hydrolase family 28 (InterPro IPR000743), were accessed from InterPro under the UniProt identifiers listed above. A percent identity matrix was generated by Clustal2.1 from the PGRA1 and PGRA2 full protein sequence alignment created by MUSCLE (version 3.8.425). AlphaFold models for AT1G05650 (Q9SYK6) and AT1G05660 (Q9SYK7) were obtained from AlphaFoldDB. Structural alignments were performed in UCSF ChimeraX (version 1.9; (Goddard et al., 2018)) using Matchmaker tool to align the AlphaFold models of AT1G05650 and AT1G05660 to known structures used as template structures: Arabidopsis Endo-PG (PDB: 7B7A) and Yersinia enterocolitica Exo-PG (PDB: 2UVE).

### Phylogenetic analysis

Evolutionary analyses were conducted in MEGA11 (version 11.0.13; (Tamura et al., 2021)). For broad phylogenetic analysis, protein sequences from 25 Endo-PG and 52 Exo-PG reviewed accessions were retrieved from UniProt (search conducted on 09-01-2025), alongside PGRA1 and PGRA2 protein sequences. Sequences were imported into MEGA11 software and signal peptide was removed (23 amino acid residues from the N-terminus) from each protein sequence before performing an initial MSA using MUSCLE. From the initial MSA, variable regions were further trimmed (49 MSA positions from the N-terminus and 28 MSA positions from the C-terminus) and re-aligned. This alignment was used to infer evolutionary history by using the Maximum Likelihood method and JTT matrix-based model (initial tree obtained by Neighbor-Joining method). The bootstrap consensus tree was inferred from 100 replicates. Branches corresponding to partitions reproduced in less than 50% bootstrap replicates are collapsed. The plant PG phylogenetic analysis was done analogously, only with 45 and 15 MSA positions trimmed from the N- and C-termini, respectively, and with bootstrap consensus tree inferred from 1000 replicates.

### Statistical and graphical analysis

Figures were assembled in Inkscape. Individual dots in boxplots represent individual data points, the number of individual measurements is specified; whiskers extend to data points that are less than 1.5 x interquartile range away from 1st/3rd quartile. Line graphs were made by Microsoft Excel. Error bars are SD (Standard Deviation) in line graphs.

All experiments using lines prepared in this study were conducted with at least three independent homozygous lines. Each experiment was repeated in at least three biological replicates, and representative results are shown here. The statistical test used is indicated for each figure.

## References

Abbott, D. W., & Boraston, A. B. (2007). The structural basis for exopolygalacturonase activity in a family 28 glycoside hydrolase. Journal of Molecular Biology, 368(5), 1215– 1222.

Anderson, C. T. (2019). Pectic polysaccharides in plants: Structure, biosynthesis, functions, and applications. In Biologically-Inspired Systems (pp. 487–514). Springer International Publishing.

Bao, D., Chang, S., Li, X., & Qi, Y. (2024). Advances in the study of auxin early response genes: Aux/IAA, GH3, and SAUR. The Crop Journal, 12(4), 964–978.

Barbez, E., Dünser, K., Gaidora, A., Lendl, T., & Busch, W. (2017). Auxin steers root cell expansion via apoplastic pH regulation in Arabidopsis thaliana. Proceedings of the National Academy of Sciences of the United States of America, 114(24), E4884–E4893.

Barnes, W. J., Zelinsky, E., & Anderson, C. T. (2022). Polygalacturonase activity promotes aberrant cell separation in the quasimodo2 mutant of Arabidopsis thaliana. Cell Surface (Amsterdam), 8(100069), 100069.

Bindels, D. S., Haarbosch, L., van Weeren, L., Postma, M., Wiese, K. E., Mastop, M., Aumonier, S., Gotthard, G., Royant, A., Hink, M. A., & Gadella, T. W. J., Jr. (2017). mScarlet: a bright monomeric red fluorescent protein for cellular imaging. Nature Methods, 14(1), 53–56.

Braybrook, S. A., & Peaucelle, A. (2013). Mechano-chemical aspects of organ formation in Arabidopsis thaliana: the relationship between auxin and pectin. PloS One, 8(3), e57813.

Cancé, C., Martin-Arevalillo, R., Boubekeur, K., & Dumas, R. (2022). Auxin response factors are keys to the many auxin doors. The New Phytologist, 235(2), 402–419.

Cao, L., Lu, W., Mata, A., Nishinari, K., & Fang, Y. (2020). Egg-box model-based gelation of alginate and pectin: A review. Carbohydrate Polymers, 242(116389), 116389.

Capodicasa, C., Vairo, D., Zabotina, O., McCartney, L., Caprari, C., Mattei, B., Manfredini, C., Aracri, B., Benen, J., Knox, J. P., De Lorenzo, G., & Cervone, F. (2004). Targeted modification of homogalacturonan by transgenic expression of a fungal polygalacturonase alters plant growth. Plant Physiology, 135(3), 1294–1304.

Chebli, Y., & Geitmann, A. (2017). Cellular growth in plants requires regulation of cell wall biochemistry. Current Opinion in Cell Biology, 44, 28–35.

Cosgrove, D. J. (1993). How do plant cell walls extend? Plant Physiology, 102(1), 1–6.

Cosgrove, Daniel J. (2018a). Diffuse growth of plant cell walls. Plant Physiology, 176(1), 16–27.

Cosgrove, Daniel J. (2018b). Nanoscale structure, mechanics and growth of epidermal cell walls. Current Opinion in Plant Biology, 46, 77–86.

Cosgrove, Daniel J. (2022). Building an extensible cell wall. Plant Physiology, 189(3), 1246–1277.

Dash, L., Swaminathan, S., Šimura, J., Gonzales, C. L. P., Montes, C., Solanki, N., Mejia, L., Ljung, K., Zabotina, O. A., & Kelley, D. R. (2023). Changes in cell wall composition due to a pectin biosynthesis enzyme GAUT10 impact root growth. Plant Physiology, 193(4), 2480–2497.

Dauphin, B. G., Ranocha, P., Dunand, C., & Burlat, V. (2022). Cell-wall microdomain remodeling controls crucial developmental processes. Trends in Plant Science, 27(10), 1033–1048.

Delmer, D., Dixon, R. A., Keegstra, K., & Mohnen, D. (2024). The plant cell wall-dynamic, strong, and adaptable-is a natural shapeshifter. The Plant Cell, 36(5), 1257–1311.

Duan, Q., Liu, M.-C. J., Kita, D., Jordan, S. S., Yeh, F.-L. J., Yvon, R., Carpenter, H., Federico, A. N., Garcia-Valencia, L. E., Eyles, S. J., Wang, C.-S., Wu, H.-M., & Cheung, A. Y. (2020). FERONIA controls pectin- and nitric oxide-mediated male-female interaction. Nature, 579(7800), 561–566.

Dubey, S. M., Han, S., Stutzman, N., Prigge, M. J., Medvecká, E., Platre, M. P., Busch, W., Fendrych, M., & Estelle, M. (2023). The AFB1 auxin receptor controls the cytoplasmic auxin response pathway in Arabidopsis thaliana. Molecular Plant, 16(7), 1120–1130.

Dünser, K., Gupta, S., Herger, A., Feraru, M. I., Ringli, C., & Kleine-Vehn, J. (2019). Extracellular matrix sensing by FERONIA and Leucine-Rich Repeat Extensins controls vacuolar expansion during cellular elongation in Arabidopsis thaliana. The EMBO Journal, 38(7), e100353.

Dusek, J., Plchova, H., Cerovska, N., Poborilova, Z., Navratil, O., Kratochvilova, K., Gunter, C., Jacobs, R., Hitzeroth, I. I., Rybicki, E. P., & Moravec, T. (2020). Extended set of GoldenBraid compatible vectors for fast assembly of multigenic constructs and their use to create geminiviral expression vectors. Frontiers in Plant Science, 11. 10.3389/fpls.2020.522059

Duvetter, T., Fraeye, I., Sila, D. N., Verlent, I., Smout, C., Hendrickx, M., & Van Loey, A. (2006). Mode of de-esterification of alkaline and acidic pectin methyl esterases at different pH conditions. Journal of Agricultural and Food Chemistry, 54(20), 7825–7831.

Ferrari, S. (2013). Oligogalacturonides: plant damage-associated molecular patterns and regulators of growth and development. Frontiers in Plant Science, 4. 10.3389/fpls.2013.00049

Friml, J., Wiśniewska, J., Benková, E., Mendgen, K., & Palme, K. (2002). Lateral relocation of auxin efflux regulator PIN3 mediates tropism in Arabidopsis. Nature, 415(6873), 806– 809.

Fry, S. C. (1988). The growing plant cell wall. John Wiley & Sons.

Gallemí, M., Montesinos, J. C., Zarevski, N., Pribyl, J., Skládal, P., Hannezo, E., & Benková, E. (2022). Dual role of Pectin Methyl Esterase activity in the regulation of plant cell wall biophysical properties. In bioRxiv. 10.1101/2022.06.14.495617

Goddard, T. D., Huang, C. C., Meng, E. C., Pettersen, E. F., Couch, G. S., Morris, J. H., & Ferrin, T. E. (2018). UCSF ChimeraX: Meeting modern challenges in visualization and analysis. Protein Science: A Publication of the Protein Society, 27(1), 14–25.

Gonzalez, J. H., Taylor, J. S., Reed, K. M., Wright, R. C., & Bargmann, B. O. R. (2021). Temporal control of morphogenic factor expression determines efficacy in enhancing regeneration. Plants, 10(11), 2271.

González-Carranza, Z. H., Elliott, K. A., & Roberts, J. A. (2007). Expression of polygalacturonases and evidence to support their role during cell separation processes in Arabidopsis thaliana. Journal of Experimental Botany, 58(13), 3719– 3730.

Goubet, F., Jackson, P., Deery, M. J., & Dupree, P. (2002). Polysaccharide analysis using carbohydrate gel electrophoresis: a method to study plant cell wall polysaccharides and polysaccharide hydrolases. Analytical Biochemistry, 300(1), 53–68.

Gould, S. J., & Subramani, S. (1988). Firefly luciferase as a tool in molecular and cell biology. Analytical Biochemistry, 175(1), 5–13.

Haas, K. T., Wightman, R., Meyerowitz, E. M., & Peaucelle, A. (2020). Pectin homogalacturonan nanofilament expansion drives morphogenesis in plant epidermal cells. Science (New York, N.Y.), 367(6481), 1003–1007.

Haas, K. T., Wightman, R., Peaucelle, A., & Höfte, H. (2021). The role of pectin phase separation in plant cell wall assembly and growth. Cell Surface (Amsterdam), 7(100054), 100054.

Hager, A., Menzel, H., & Krauss, A. (1971). Experiments and hypothesis concerning the primary action of auxin in elongation growth. Planta, 100(1), 47–75.

He, W., Brumos, J., Li, H., Ji, Y., Ke, M., Gong, X., Zeng, Q., Li, W., Zhang, X., An, F., Wen, X., Li, P., Chu, J., Sun, X., Yan, C., Yan, N., Xie, D.-Y., Raikhel, N., Yang, Z., … Guo, H. (2011). A small-molecule screen identifies L-kynurenine as a competitive inhibitor of TAA1/TAR activity in ethylene-directed auxin biosynthesis and root growth in Arabidopsis. The Plant Cell, 23(11), 3944–3960.

Henrissat, B., & Davies, G. (1997). Structural and sequence-based classification of glycoside hydrolases. Current Opinion in Structural Biology, 7(5), 637–644.

Hewezi, T., Howe, P., Maier, T. R., Hussey, R. S., Mitchum, M. G., Davis, E. L., & Baum, T. J. (2008). Cellulose binding protein from the parasitic nematode Heterodera schachtii interacts with Arabidopsis pectin methylesterase: cooperative cell wall modification during parasitism. The Plant Cell, 20(11), 3080–3093.

Hocq, L., Guinand, S., Habrylo, O., Voxeur, A., Tabi, W., Safran, J., Fournet, F., Domon, J.-M., Mollet, J.-C., Pilard, S., Pau-Roblot, C., Lehner, A., Pelloux, J., & Lefebvre, V. (2020). The exogenous application of AtPGLR, an endo-polygalacturonase, triggers pollen tube burst and repair. The Plant Journal: For Cell and Molecular Biology, 103(2), 617–633.

Hocq, L., Habrylo, O., Sénéchal, F., Voxeur, A., Pau-Roblot, C., Safran, J., Fournet, F., Bassard, S., Battu, V., Demailly, H., Tovar, J. C., Pilard, S., Marcelo, P., Savary, B. J., Mercadante, D., Njo, M. F., Beeckman, T., Boudaoud, A., Gutierrez, L., … Lefebvre, V. (2024). Mutation of AtPME2, a pH-dependent pectin methylesterase, affects cell wall structure and hypocotyl elongation. Plant & Cell Physiology, 65(2), 301–318.

Hocq, L., Pelloux, J., & Lefebvre, V. (2017). Connecting homogalacturonan-type pectin remodeling to acid growth. Trends in Plant Science, 22(1), 20–29.

Hocq, L., Sénéchal, F., Lefebvre, V., Lehner, A., Domon, J.-M., Mollet, J.-C., Dehors, J., Pageau, K., Marcelo, P., Guérineau, F., Kolšek, K., Mercadante, D., & Pelloux, J. (2017). Combined experimental and computational approaches reveal distinct pH dependence of pectin methylesterase inhibitors. Plant Physiology, 173(2), 1075–1093.

Huang, D. W., Sherman, B. T., & Lempicki, R. A. (2009). Systematic and integrative analysis of large gene lists using DAVID bioinformatics resources. Nature Protocols, 4(1), 44–57.

Huerta, A. I., Sancho-Andrés, G., Montesinos, J. C., Silva-Navas, J., Bassard, S., Pau-Roblot, C., Kesten, C., Schlechter, R., Dora, S., Ayupov, T., Pelloux, J., Santiago, J., & Sánchez-Rodríguez, C. (2023). The WAK-like protein RFO1 acts as a sensor of the pectin methylation status in Arabidopsis cell walls to modulate root growth and defense. Molecular Plant, 16(5), 865–881.

Jeong, H. Y., Nguyen, H. P., Eom, S. H., & Lee, C. (2018). Integrative analysis of pectin methylesterase (PME) and PME inhibitors in tomato (Solanum lycopersicum): Identification, tissue-specific expression, and biochemical characterization. Plant Physiology and Biochemistry, 132, 557–565.

Jobert, F., Soriano, A., Brottier, L., Casset, C., Divol, F., Safran, J., Lefebvre, V., Pelloux, J., Robert, S., & Péret, B. (2021). Auxin and pectin remodeling interplay during rootlet emergence in white lupin. In bioRxiv. bioRxiv. 10.1101/2021.07.19.452882

Jobert, F., Yadav, S., & Robert, S. (2023). Auxin as an architect of the pectin matrix. Journal of Experimental Botany, 74(22), 6933–6949.

Jonsson, K., Lathe, R. S., Kierzkowski, D., Routier-Kierzkowska, A.-L., Hamant, O., & Bhalerao, R. P. (2021). Mechanochemical feedback mediates tissue bending required for seedling emergence. Current Biology: CB, 31(6), 1154-1164.e3.

Kim, J., & Patterson, S. E. (2006). Expression divergence and functional redundancy of polygalacturonases in floral organ abscission. Plant Signaling & Behavior, 1(6), 281–283.

Kim, J., Shiu, S.-H., Thoma, S., Li, W.-H., & Patterson, S. E. (2006). Patterns of expansion and expression divergence in the plant polygalacturonase gene family. Genome Biology, 7(9), R87.

Kubalova, M., Griffiths, J., Muller, K., Jones, A. M., & Fendrych, M. (2025). Gibberellin-deactivating GA2OX enzymes act as a hub for auxin-gibberellin crosstalk in Arabidopsis thaliana root growth regulation. In bioRxiv. 10.1101/2025.02.03.636207

Kubalová, M., Müller, K., Dobrev, P. I., Rizza, A., Jones, A. M., & Fendrych, M. (2024). Auxin co-receptor IAA17/AXR3 controls cell elongation in Arabidopsis thaliana root solely by modulation of nuclear auxin pathway. The New Phytologist, 241(6), 2448–2463.

Kubalová, M., Schmidtová, M., & Fendrych, M. (2025). Unresolved roles of Aux/IAA proteins in auxin responses. Physiologia Plantarum, 177(2), e70221.

Leng, Y., Yang, Y., Ren, D., Huang, L., Dai, L., Wang, Y., Chen, L., Tu, Z., Gao, Y., Li, X., Zhu, L., Hu, J., Zhang, G., Gao, Z., Guo, L., Kong, Z., Lin, Y., Qian, Q., & Zeng, D. (2017). A rice PECTATE LYASE-LIKE gene is required for plant growth and leaf senescence. Plant Physiology, 174(2), 1151–1166.

Lewis, D. R., Olex, A. L., Lundy, S. R., Turkett, W. H., Fetrow, J. S., & Muday, G. K. (2013). A kinetic analysis of the auxin transcriptome reveals cell wall remodeling proteins that modulate lateral root development in Arabidopsis. The Plant Cell, 25(9), 3329–3346.

Leyser, O. (2018). Auxin signaling. Plant Physiology, 176(1), 465–479.

Li, L., Verstraeten, I., Roosjen, M., Takahashi, K., Rodriguez, L., Merrin, J., Chen, J., Shabala, L., Smet, W., Ren, H., Vanneste, S., Shabala, S., De Rybel, B., Weijers, D., Kinoshita, T., Gray, W. M., & Friml, J. (2021). Cell surface and intracellular auxin signalling for H+ fluxes in root growth. Nature, 599(7884), 273–277.

Li, Y., Smith, C., Corke, F., Zheng, L., Merali, Z., Ryden, P., Derbyshire, P., Waldron, K., & Bevan, M. W. (2007). Signaling from an altered cell wall to the nucleus mediates sugar-responsive growth and development in Arabidopsis thaliana. The Plant Cell, 19(8), 2500–2515.

Lin, W., Zhou, X., Tang, W., Takahashi, K., Pan, X., Dai, J., Ren, H., Zhu, X., Pan, S., Zheng, H., Gray, W. M., Xu, T., Kinoshita, T., & Yang, Z. (2021). TMK-based cell-surface auxin signalling activates cell-wall acidification. Nature, 599(7884), 278–282.

Lintilhac, P. M. (2014). The problem of morphogenesis: unscripted biophysical control systems in plants. Protoplasma, 251(1), 25–36.

Mähönen, A. P., Tusscher, K. T., Siligato, R., Smetana, O., Díaz-Triviño, S., Salojärvi, J., Wachsman, G., Prasad, K., Heidstra, R., & Scheres, B. (2014). PLETHORA gradient formation mechanism separates auxin responses. Nature, 515(7525), 125–129.

Majda, M., & Robert, S. (2018). The role of auxin in cell wall expansion. International Journal of Molecular Sciences, 19(4). 10.3390/ijms19040951

Mase, K., Mizuno, H., Nakamichi, N., Suzuki, T., Kojima, T., Kamiya, S., Takeuchi, T., Kondo, C., Yamashita, H., Sakaoka, S., Morikami, A., & Tsukagoshi, H. (2023). AtMYB50 regulates root cell elongation by upregulating PECTIN METHYLESTERASE INHIBITOR 8 in Arabidopsis thaliana. PloS One, 18(12), e0285241.

Molina, A., Miedes, E., Bacete, L., Rodríguez, T., Mélida, H., Denancé, N., Sánchez-Vallet, A., Rivière, M.-P., López, G., Freydier, A., Barlet, X., Pattathil, S., Hahn, M., & Goffner, D. (2021). Arabidopsis cell wall composition determines disease resistance specificity and fitness. Proceedings of the National Academy of Sciences of the United States of America, 118(5), e2010243118.

Mravec, J., Kračun, S. K., Rydahl, M. G., Westereng, B., Miart, F., Clausen, M. H., Fangel, J. U., Daugaard, M., Van Cutsem, P., De Fine Licht, H.H., Höfte, H., Malinovsky, F. G., Domozych, D. S., & Willats, W. G. T. (2014). Tracking developmentally regulated post-synthetic processing of homogalacturonan and chitin using reciprocal oligosaccharide probes. Development (Cambridge, England), 141(24), 4841–4850.

Nagayama, T., Tatsumi, A., Nakamura, A., Yamaji, N., Satoh, S., Furukawa, J., & Iwai, H. (2022). Effects of polygalacturonase overexpression on pectin distribution in the elongation zones of roots under aluminium stress. AoB Plants, 14(2), ac003.

Nemhauser, J. L., Hong, F., & Chory, J. (2006). Different plant hormones regulate similar processes through largely nonoverlapping transcriptional responses. Cell, 126(3), 467–475.

Nishitani, K., & Masuda, Y. (1981). Auxin-induced changes in the cell wall structure: Changes in the sugar compositions, intrinsic viscosity and molecular weight distributions of matrix polysaccharides of the epicotyl cell wall of Vigna angularis. Physiologia Plantarum, 52(4), 482–494.

Ogawa, M., Kay, P., Wilson, S., & Swain, S. M. (2009). ARABIDOPSIS DEHISCENCE ZONE POLYGALACTURONASE1 (ADPG1), ADPG2, and QUARTET2 are polygalacturonases required for cell separation during reproductive development inArabidopsis. The Plant Cell, 21(1), 216–233.

Overvoorde, P., Fukaki, H., & Beeckman, T. (2010). Auxin control of root development. Cold Spring Harbor Perspectives in Biology, 2(6), a001537–a001537.

Pattathil, S., Avci, U., Baldwin, D., Swennes, A. G., McGill, J. A., Popper, Z., Bootten, T., Albert, A., Davis, R. H., Chennareddy, C., Dong, R., O’Shea, B., Rossi, R., Leoff, C., Freshour, G., Narra, R., O’Neil, M., York, W. S., & Hahn, M. G. (2010). A comprehensive toolkit of plant cell wall glycan-directed monoclonal antibodies. Plant Physiology, 153(2), 514–525.

Peaucelle, A., Braybrook, S. A., Le Guillou, L., Bron, E., Kuhlemeier, C., & Höfte, H. (2011). Pectin-induced changes in cell wall mechanics underlie organ initiation in Arabidopsis. Current Biology: CB, 21(20), 1720–1726.

Phyo, P., Gu, Y., & Hong, M. (2019). Impact of acidic pH on plant cell wall polysaccharide structure and dynamics: insights into the mechanism of acid growth in plants from solid-state NMR. Cellulose (London, England), 26(1), 291–304.

Pottier, D., Roitsch, T., & Persson, S. (2023). Cell wall regulation by carbon allocation and sugar signaling. Cell Surface (Amsterdam), 9(100096), 100096.

Rayle, D. L., & Cleland, R. (1970). Enhancement of wall loosening and elongation by Acid solutions. Plant Physiology, 46(2), 250–253.

Rhee, S. Y., Osborne, E., Poindexter, P. D., & Somerville, C. R. (2003). Microspore separation in the quartet 3 mutants of Arabidopsis is impaired by a defect in a developmentally regulated polygalacturonase required for pollen mother cell wall degradation. Plant Physiology, 133(3), 1170–1180.

Rößling, A.-K., Dünser, K., Liu, C., Lauw, S., Rodriguez-Franco, M., Kalmbach, L., Barbez, E., & Kleine-Vehn, J. (2024). Pectin methylesterase activity is required for RALF1 peptide signalling output. 10.7554/elife.96943.2

Roychoudhry, S., & Kepinski, S. (2022). Auxin in root development. Cold Spring Harbor Perspectives in Biology, 14(4), a039933.

Rui, Y., Xiao, C., Yi, H., Kandemir, B., Wang, J. Z., Puri, V. M., & Anderson, C. T. (2017). POLYGALACTURONASE INVOLVED IN EXPANSION3 functions in seedling development, rosette growth, and stomatal dynamics in Arabidopsis thaliana. The Plant Cell, 29(10), 2413–2432.

Safran, J., Tabi, W., Ung, V., Lemaire, A., Habrylo, O., Bouckaert, J., Rouffle, M., Voxeur, A., Pongrac, P., Bassard, S., Molinié, R., Fontaine, J.-X., Pilard, S., Pau-Roblot, C., Bonnin, E., Larsen, D. S., Morel-Rouhier, M., Girardet, J.-M., Lefebvre, V., … Pelloux, J. (2023). Plant polygalacturonase structures specify enzyme dynamics and processivities to fine-tune cell wall pectins. The Plant Cell, 35(8), 3073– 3091.

Samalova, M., Fricker, M., & Moore, I. (2006). Ratiometric fluorescence-imaging assays of plant membrane traffic using polyproteins. Traffic (Copenhagen, Denmark), 7(12), 1701–1723.

Santiago-Doménech, N., Jiménez-Bemúdez, S., Matas, A. J., Rose, J. K. C., Muñoz-Blanco, J., Mercado, J. A., & Quesada, M. A. (2008). Antisense inhibition of a pectate lyase gene supports a role for pectin depolymerization in strawberry fruit softening. Journal of Experimental Botany, 59(10), 2769–2779.

Sarrion-Perdigones, A., Vazquez-Vilar, M., Palaci, J., Castelijns, B., Forment, J., Ziarsolo, P., Blanca, J., Granell, A., & Orzaez, D. (2013). GoldenBraid 2.0: A comprehensive DNA assembly framework for plant synthetic biology. Plant Physiology, 162(3), 1618–1631.

Sarrion-Perdigones, Alejandro, Falconi, E. E., Zandalinas, S. I., Juárez, P., Fernández-del-Carmen, A., Granell, A., & Orzaez, D. (2011). GoldenBraid: an iterative cloning system for standardized assembly of reusable genetic modules. PloS One, 6(7), e21622.

Sénéchal, F., Graff, L., Surcouf, O., Marcelo, P., Rayon, C., Bouton, S., Mareck, A., Mouille, G., Stintzi, A., Höfte, H., Lerouge, P., Schaller, A., & Pelloux, J. (2014). Arabidopsis PECTIN METHYLESTERASE17 is co-expressed with and processed by SBT3.5, a subtilisin-like serine protease. Annals of Botany, 114(6), 1161–1175.

Serre, N. B. C., & Fendrych, M. (2022). ACORBA: Automated workflow to measure Arabidopsis thaliana root tip angle dynamics. Quantitative Plant Biology, 3(e9), e9.

Serre, N. B. C., Kralík, D., Yun, P., Slouka, Z., Shabala, S., & Fendrych, M. (2021). AFB1 controls rapid auxin signalling through membrane depolarization in Arabidopsis thaliana root. Nature Plants, 7(9), 1229–1238.

Serre, N. B. C., Wernerová, D., Vittal, P., Dubey, S. M., Medvecká, E., Jelínková, A., Petrášek, J., Grossmann, G., & Fendrych, M. (2023). The AUX1-AFB1-CNGC14 module establishes a longitudinal root surface pH profile. ELife, 12. 10.7554/elife.85193

Shaner, N. C., Campbell, R. E., Steinbach, P. A., Giepmans, B. N. G., Palmer, A. E., & Tsien, R. Y. (2004). Improved monomeric red, orange and yellow fluorescent proteins derived from Discosoma sp. red fluorescent protein. Nature Biotechnology, 22(12), 1567–1572.

Sun, H., Hao, P., Gu, L., Cheng, S., Wang, H., Wu, A., Ma, L., Wei, H., & Yu, S. (2020). Pectate lyase-like Gene GhPEL76 regulates organ elongation in Arabidopsis and fiber elongation in cotton. Plant Science: An International Journal of Experimental Plant Biology, 293(110395), 110395.

Tamura, K., Stecher, G., & Kumar, S. (2021). MEGA11: Molecular Evolutionary Genetics Analysis version 11. Molecular Biology and Evolution, 38(7), 3022–3027.

Tiwari, S. B., Hagen, G., & Guilfoyle, T. (2003). The roles of auxin response factor domains in auxin-responsive transcription. The Plant Cell, 15(2), 533–543.

Verhage, L. (2021). Get in shape - how a polygalacturonase affects plant morphology. The Plant Journal: For Cell and Molecular Biology, 106(6), 1491–1492.

Verhertbruggen, Y., Marcus, S. E., Haeger, A., Ordaz-Ortiz, J. J., & Knox, J. P. (2009). An extended set of monoclonal antibodies to pectic homogalacturonan. Carbohydrate Research, 344(14), 1858–1862.

Vincken, J.-P., Schols, H. A., Oomen, R. J. F. J., McCann, M. C., Ulvskov, P., Voragen, A. G. J., & Visser, R. G. F. (2003). If homogalacturonan were a side chain of rhamnogalacturonan I. Implications for cell wall architecture. Plant Physiology, 132(4), 1781–1789.

Vogel, J. P., Raab, T. K., Schiff, C., & Somerville, S. C. (2002). PMR6, a pectate lyase–like gene required for powdery mildew susceptibility in Arabidopsis. The Plant Cell, 14(9), 2095– 2106.

von Wangenheim, D., Hauschild, R., Fendrych, M., Barone, V., Benková, E., & Friml, J. (2017). Live tracking of moving samples in confocal microscopy for vertically grown roots. ELife, 6. 10.7554/elife.26792

Voragen, A. G. J., Coenen, G.-J., Verhoef, R. P., & Schols, H. A. (2009). Pectin, a versatile polysaccharide present in plant cell walls. Structural Chemistry, 20(2), 263–275.

Wachsman, G., Zhang, J., Moreno-Risueno, M. A., Anderson, C. T., & Benfey, P. N. (2020). Cell wall remodeling and vesicle trafficking mediate the root clock in Arabidopsis. Science (New York, N.Y.), 370(6518), 819–823.

Willats, W. G., Orfila, C., Limberg, G., Buchholt, H. C., van Alebeek, G. J., Voragen, A. G., Marcus, S. E., Christensen, T. M., Mikkelsen, J. D., Murray, B. S., & Knox, J. P. (2001). Modulation of the degree and pattern of methyl-esterification of pectic homogalacturonan in plant cell walls. Implications for pectin methyl esterase action, matrix properties, and cell adhesion. The Journal of Biological Chemistry, 276(22), 19404–19413.

Wolf, S., Mouille, G., & Pelloux, J. (2009). Homogalacturonan methyl-esterification and plant development. Molecular Plant, 2(5), 851–860.

Wolf, S., Mravec, J., Greiner, S., Mouille, G., & Höfte, H. (2012). Plant cell wall homeostasis is mediated by brassinosteroid feedback signaling. Current Biology: CB, 22(18), 1732– 1737.

Wormit, A., & Usadel, B. (2018). The multifaceted role of pectin methylesterase inhibitors (PMEIs). International Journal of Molecular Sciences, 19(10), 2878.

Xiao, C., Barnes, W. J., Zamil, M. S., Yi, H., Puri, V. M., & Anderson, C. T. (2017). Activation tagging of Arabidopsis POLYGALACTURONASE INVOLVED IN EXPANSION2 promotes hypocotyl elongation, leaf expansion, stem lignification, mechanical stiffening, and lodging. The Plant Journal: For Cell and Molecular Biology, 89(6), 1159–1173.

Xiao, C., Somerville, C., & Anderson, C. T. (2014). POLYGALACTURONASE INVOLVED IN EXPANSION1 functions in cell elongation and flower development in Arabidopsis. The Plant Cell, 26(3), 1018–1035.

Xing, H.-L., Dong, L., Wang, Z.-P., Zhang, H.-Y., Han, C.-Y., Liu, B., Wang, X.-C., & Chen, Q.-J. (2014). A CRISPR/Cas9 toolkit for multiplex genome editing in plants. BMC Plant Biology, 14(1), 327.

Xu, F., Gonneau, M., Faucher, E., Habrylo, O., Lefebvre, V., Domon, J.-M., Martin, M., Sénéchal, F., Peaucelle, A., Pelloux, J., & Höfte, H. (2022). Biochemical characterization of Pectin Methylesterase Inhibitor 3 from Arabidopsis thaliana. Cell Surface (Amsterdam), 8(100080), 100080.

Yang, Y., Anderson, C. T., & Cao, J. (2021). Polygalacturonase45 cleaves pectin and links cell proliferation and morphogenesis to leaf curvature in Arabidopsis thaliana. The Plant Journal: For Cell and Molecular Biology, 106(6), 1493–1508.

Zablackis, E., Huang, J., Müller, B., Darvill, A. G., & Albersheim, P. (1995). Characterization of the cell-wall polysaccharides of Arabidopsis thaliana leaves. Plant Physiology, 107(4), 1129–1138.

Zamil, M. S., & Geitmann, A. (2017). The middle lamella-more than a glue. Physical Biology, 14(1), 015004.

Zhang, Z., Pierce, M. L., & Mort, A. J. (2007). Changes in homogalacturonans and enzymes degrading them during cotton cotyledon expansion. Phytochemistry, 68(8), 1094–1103.

Zuo, J., Niu, Q.-W., & Chua, N.-H. (2000). An estrogen receptor-based transactivator XVE mediates highly inducible gene expression in transgenic plants. The Plant Journal: For Cell and Molecular Biology, 24(2), 265–273.

