## supplemental information for "POLYGALACTURONASES REGULATED BY AUXIN facilitate root cell elongation in *Arabidopsis thaliana* via pectin remodeling"

**A**

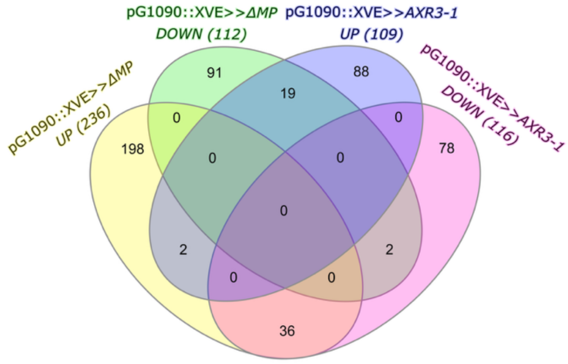


**B**

| GO term changed in pG1090::XVE>>ΔMP | | Fold enrichment | p-value |
|---|---|---|---|
| GO:0010279~indole-3-acetic acid amido synthetase activity | | 82.09812 | 1.02E-05 |
| GO:0045544~gibberellin 20-oxidase activity | | 49.25887 | 0.03978 |
| GO:1900036~positive regulation of cellular response to heat | | 42.34254 | 0.046068 |
| GO:0140964~intracellular auxin homeostasis | | 26.74266 | 4.17E-04 |
| GO:0009686~gibberellin biosynthetic process | | 18.81891 | 0.001202 |
| GO:0016762~xyloglucan:xyloglucosyl transferase activity | | 18.65866 | 1.42E-04 |
| GO:0010411~xyloglucan metabolic process | | 15.12234 | 3.22E-04 |
| GO:0048544~recognition of pollen | | 12.70276 | 0.003769 |
| GO:0009664~plant-type cell wall organization | | 11.72563 | 1.57E-04 |
| GO:0005199~structural constituent of cell wall | | 11.54505 | 0.027532 |
| GO:0009266~response to temperature stimulus | | 10.3696 | 0.006676 |
| GO:0009828~plant-type cell wall loosening | | 9.771356 | 0.037376 |
| GO:0009653~anatomical structure morphogenesis | | 9.527072 | 0.039145 |
| GO:0042546~cell wall biogenesis | | 9.479673 | 0.001896 |
| GO:0009733~response to auxin | | 9.18272 | 1.92E-15 |
| GO:0042631~cellular response to water deprivation | | 8.862392 | 0.044636 |
| GO:0030247~polysaccharide binding | | 8.396399 | 7.51E-04 |
| GO:0048367~shoot system development | | 8.28441 | 7.89E-04 |
| GO:0009826~unidimensional cell growth | | 7.757412 | 7.91E-05 |
| GO:0009734~auxin-activated signaling pathway | | 6.854009 | 4.39E-08 |
| GO:0004553~hydrolase activity, hydrolyzing O-glycosyl compounds | | 5.754541 | 0.011235 |
| GO:0009739~response to gibberellin | | 5.709107 | 0.032825 |
| GO:0009860~pollen tube growth | | 3.944957 | 0.038056 |
| GO:0048364~root development | | 3.917151 | 0.009144 |
| GO:0009738~abscisic acid-activated signaling pathway | | 3.556773 | 0.014123 |
| GO:0016491~oxidoreductase activity | | 2.981772 | 0.006733 |
| GO:0071555~cell wall organization | | 2.868366 | 0.035633 |
| GO:0000976~transcription cis-regulatory region binding | | 2.043937 | 0.033945 |
| GO:0005576~extracellular region | | -1.87859 | 0.003188 |
| GO:0005506~iron ion binding | | -3.6507 | 0.047589 |
| GO:0005975~carbohydrate metabolic process | | -4.02056 | 0.034758 |
| GO:0016829~lyase activity | | -8.77669 | 0.002523 |
| GO:0035251~UDP-glucosyltransferase activity | | -12.4006 | 0.024098 |
| GO:0004650~polygalacturonase activity | | -14.4383 | 0.002656 |
| GO:0016717~oxidoreductase activity, acting on paired donors | | -39.4276 | 0.049129 |
| GO:0016763~pentosyltransferase activity | | -46.5962 | 0.04173 |

**C**

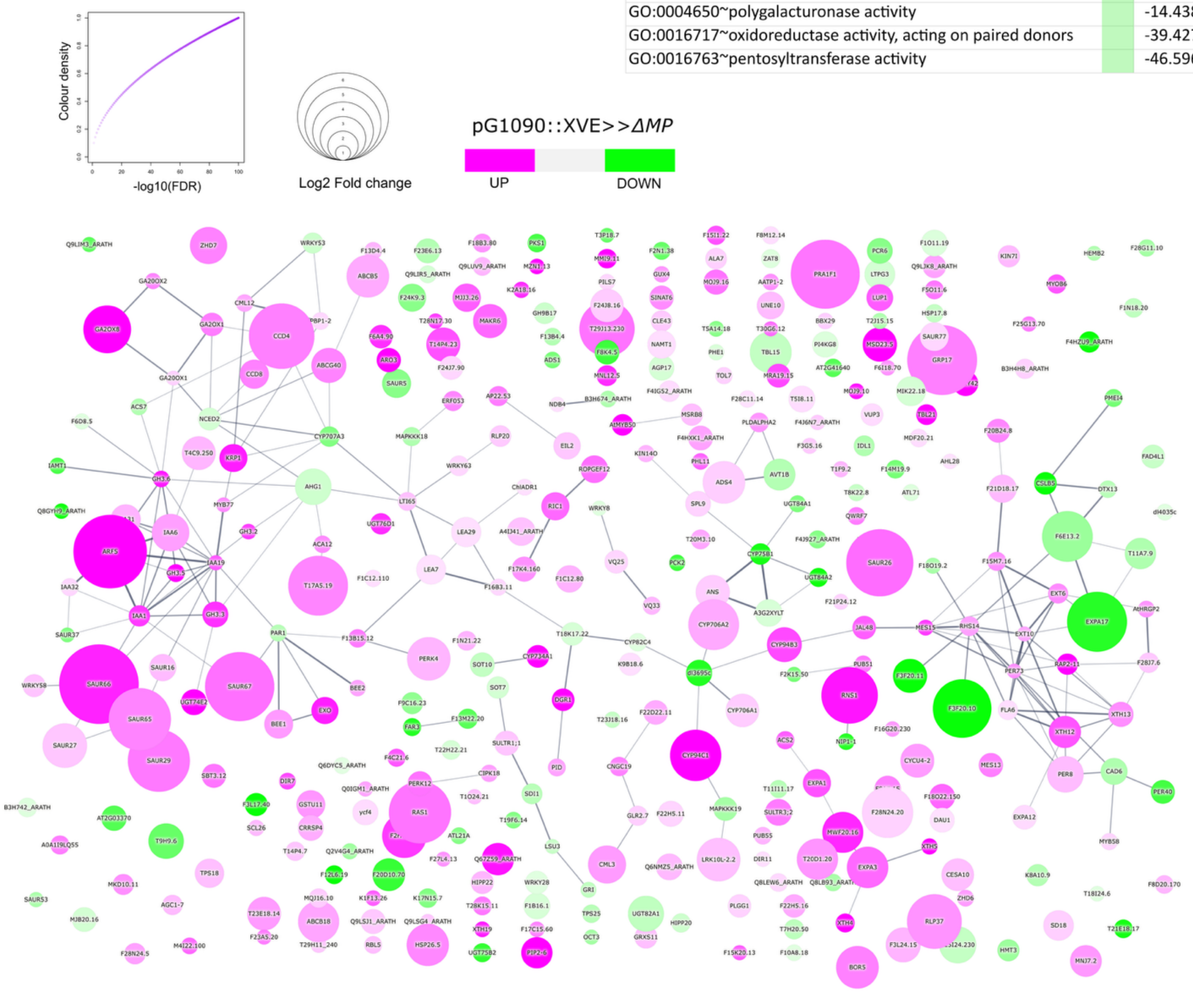

**Figure S1**

A) Venn diagram representing differentially expressed genes after 180-minute estradiol induction vs. 90-minute DMSO control treatment in pG1090::XVE>>*ΔMP* line and pG1090::XVE>>*AXR3-1* line (*FDR* ≤ 0.05 and |*fold change*| ≥ 2).

B) Gene ontology (GO) terms of differentially expressed genes after 180 min of estradiol treatment compared to 90 min of dmso treatment in pG1090::XVE>>*ΔMP* line. p-value ≤ 0.05 and Fold change ≥ 2.

C) STRING network of differentially expressed genes after 180-minute estradiol induction vs. 90-minute DMSO control treatment in pG1090::XVE>>*ΔMP* line (*FDR* ≤ 0.05 and |*fold change*| ≥ 2). Color coding indicates significant up- or downregulation, circle diameter and color intensity represent *log2FC* and √-log10(*FDR*), respectively. Connection lines indicate the confidence of all interaction types including text-mining. Complete set of DEGs is shown.

**A**

| GO term changed in pG1090::XVE>>AXR3-1 | Fold enrichment | p-value |
|---|---|---|
| GO:0103075~indole-3-pyruvate monooxygenase activity | 96.6763 | 8.12E-06 |
| GO:0033198~response to ATP | 87.58857 | 2.23E-02 |
| GO:0004499~N,N-dimethylaniline monooxygenase activity | 56.96996 | 6.00E-08 |
| GO:0010262~somatic embryogenesis | 41.80364 | 2.26E-03 |
| GO:0009851~auxin biosynthetic process | 33.14162 | 2.30E-04 |
| GO:0050661~NADP binding | 22.46703 | 6.99E-06 |
| GO:0050660~flavin adenine dinucleotide binding | 18.76658 | 1.69E-05 |
| GO:0004650~polygalacturonase activity | 11.23351 | 0.028929 |
| GO:0140825~lactoperoxidase activity | 10.09594 | 0.035188 |
| GO:0042744~hydrogen peroxide catabolic process | 9.481237 | 3.91E-02 |
| GO:0006869~lipid transport | 9.016471 | 4.28E-02 |
| GO:0071456~cellular response to hypoxia | 6.106773 | 8.78E-03 |
| GO:0009505~plant-type cell wall | 4.164497 | 0.003058 |
| GO:0020037~heme binding | 4.069283 | 1.58E-02 |
| GO:0005576~extracellular region | 2.011227 | 9.61E-04 |
| GO:0004674~protein serine/threonine kinase activity | -2.77727 | 0.009702 |
| GO:0106310~protein serine kinase activity | -2.81069 | 0.02342 |
| GO:0006979~response to oxidative stress | -5.29404 | 0.005346 |
| GO:0048364~root development | -5.88874 | 0.010047 |
| GO:0009826~unidimensional cell growth | -6.12249 | 0.08475 |
| GO:0009860~pollen tube growth | -6.64221 | 0.021911 |
| GO:0042744~hydrogen peroxide catabolic process | -8.26852 | 0.05017 |
| GO:0030247~polysaccharide binding | -8.36021 | 0.04952 |
| GO:0004553~hydrolase activity, hydrolyzing O-glycosyl compounds | -9.16758 | 0.009418 |
| GO:0140825~lactoperoxidase activity | -9.31264 | 0.040795 |
| GO:0042546~cell wall biogenesis | -11.9708 | 0.025569 |
| GO:0048367~shoot system development | -14.5298 | 3.79E-04 |
| GO:0010411~xyloglucan metabolic process | -19.0963 | 0.010566 |
| GO:0016762~xyloglucan:xyloglucosyl transferase activity | -22.2939 | 0.007882 |
| GO:0051723~protein methylesterase activity | -23.3555 | 0.081577 |
| GO:0048766~root hair initiation | -25.4618 | 0.074863 |
| GO:0009664~plant-type cell wall organization | -28.7914 | 1.48E-07 |
| GO:0005199~structural constituent of cell wall | -38.3176 | 8.35E-06 |

**B**

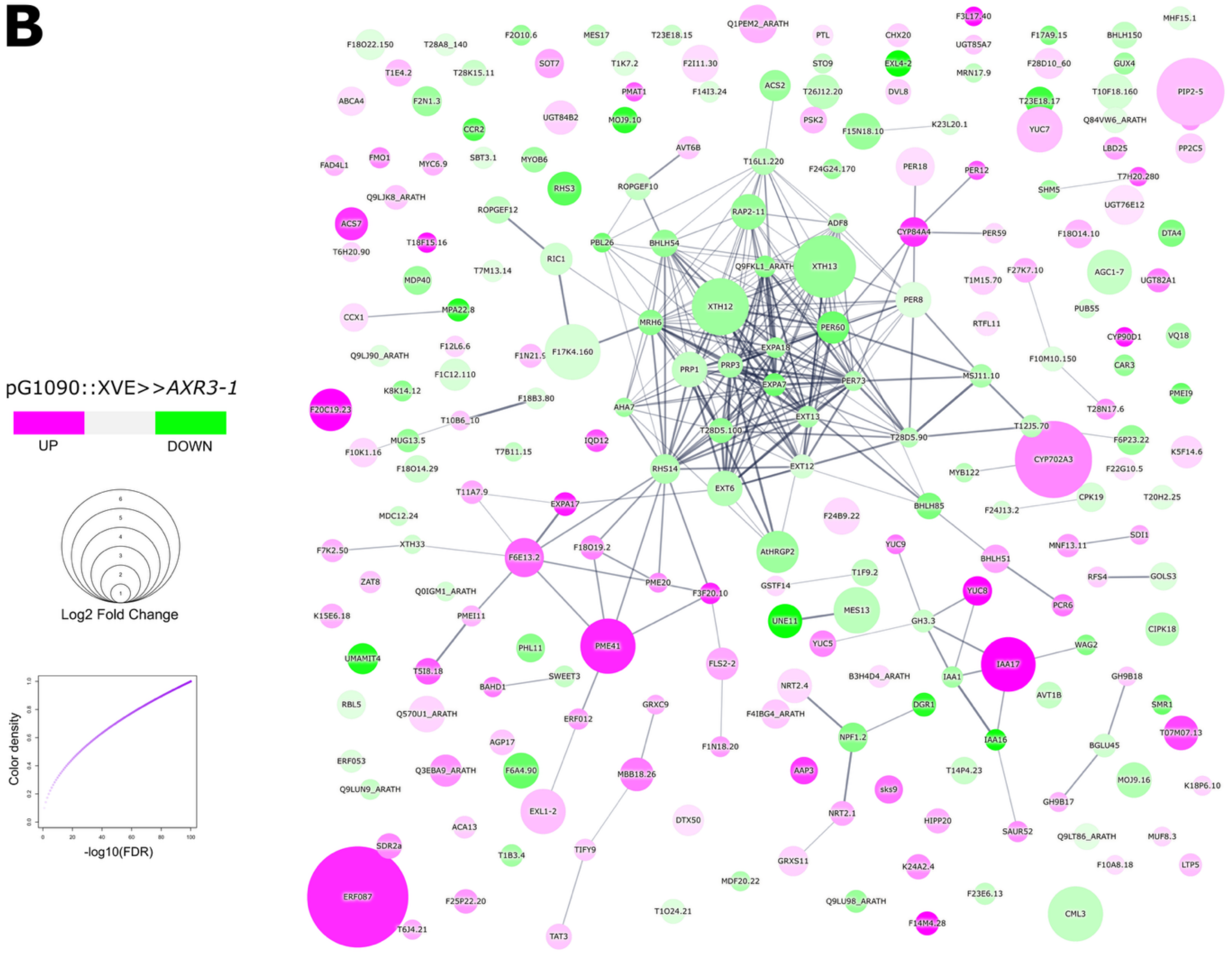

**Figure S2**

A) Gene ontology (GO) terms of differentially expressed genes after 180 min of estradiol treatment compared to 90 min of dmso treatment in pG1090::XVE>>*AXR3-1* line. p-value ≤ 0.05 and Fold change ≥ 2.

B) STRING network of differentially expressed genes after 180-minute estradiol induction vs. 90-minute DMSO control treatment in pG1090::XVE>>*AXR3-1* line (*FDR* ≤ 0.05 and |*fold change*| ≥ 2). Color coding indicates significant up- or downregulation, circle diameter and color intensity represent *log2FC* and √-log10(*FDR*), respectively. Connection lines indicate the confidence of all interaction types including text-mining. Complete set of differentially expressed genes is shown.

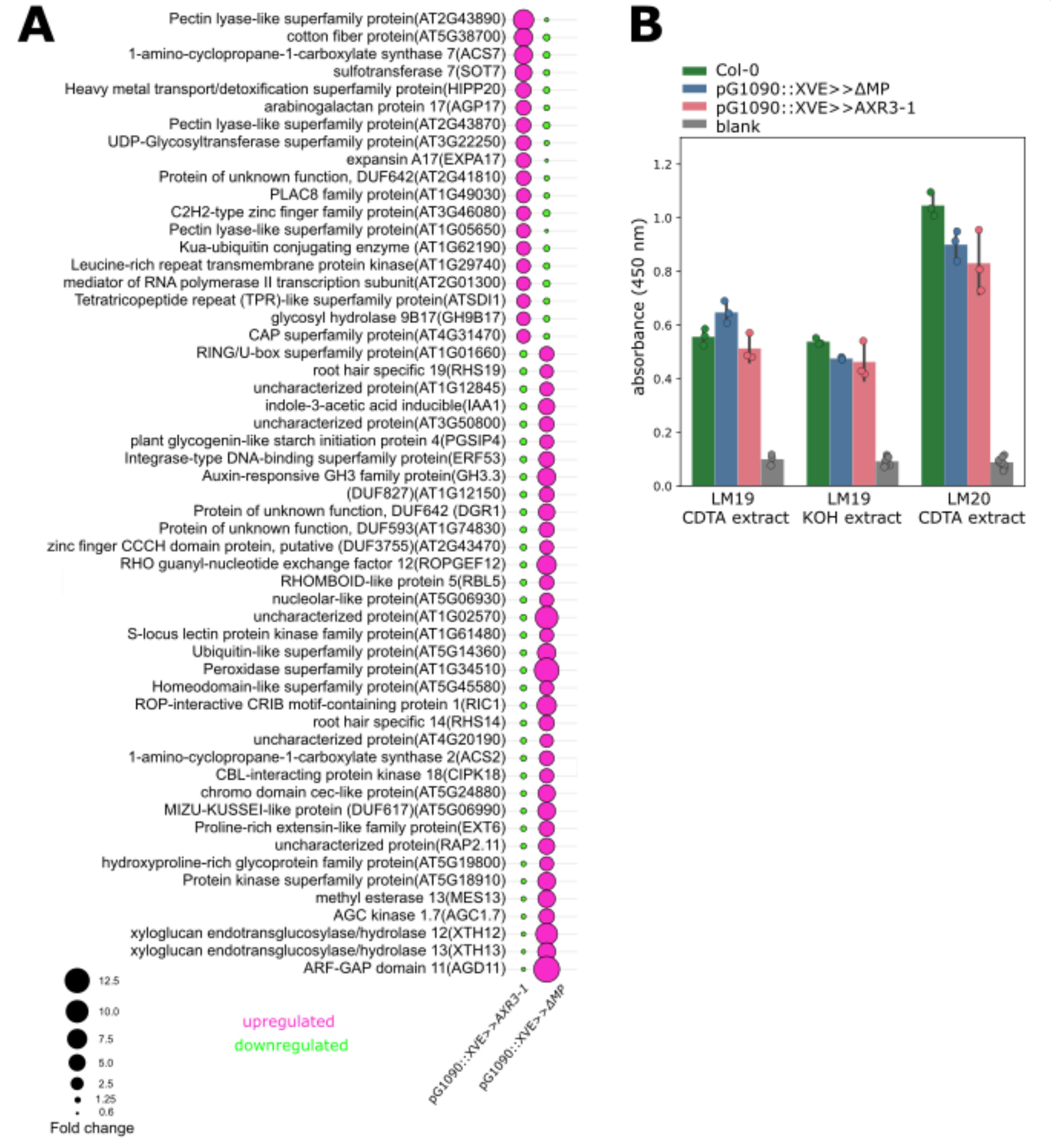


**Figure S3**

A) Full set of differentially expressed genes after 180 min of estradiol treatment compared to 90 min of dmso treatment in pG1090::XVE>>*ΔMP* and pG1090::XVE>>*AXR3-1* lines. The circle diameter indicates the fold change, with magenta representing upregulation and green representing downregulation genes.

B) Quantification of pectic epitopes in CDTA and KOH cell wall extracts of Col-0, pG1090::XVE>>*ΔMP* and pG1090::XVE>>*AXR3-1* roots probed with LM19 and LM20 monoclonal antibodies, which specifically bind to demethylesterified and methylesterified homogalacturonan epitopes, respectively.

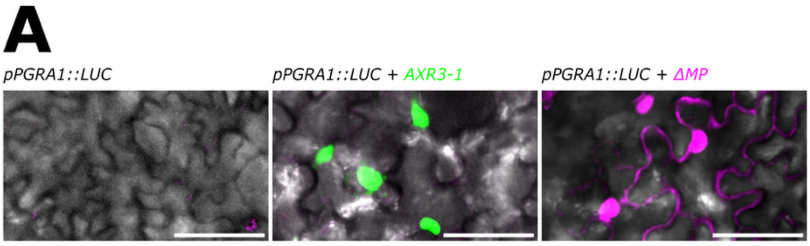


**Figure S4**
A) Tobacco leaf cells co-infiltrated with *pPGRA1::LUC* and p35S::*ΔMP*-mScarlet or p35S::*AXR3-1-mVenus*. Scale bar = 50µm.

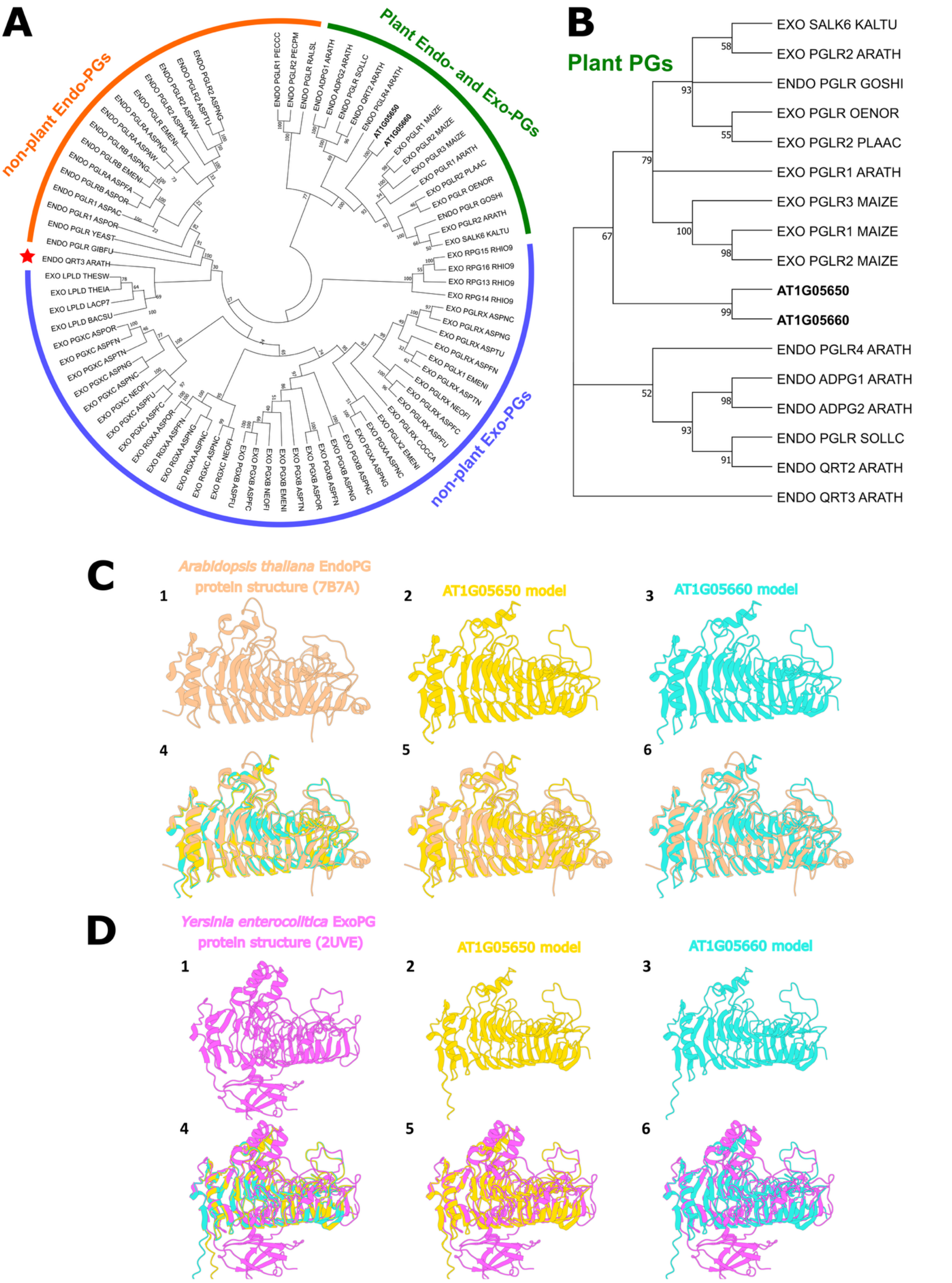
A
non-plant Endo-PGs
Plant Endo- and Exo-PGs
non-plant Exo-PGs
B
Plant PGs
EXO SALK6 KALTU
EXO PGLR2 ARATH
ENDO PGLR GOSHI
EXO PGLR OENOR
EXO PGLR2 PLAAC
EXO PGLR1 ARATH
EXO PGLR3 MAIZE
EXO PGLR1 MAIZE
EXO PGLR2 MAIZE
AT1G05650
AT1G05660
ENDO PGLR4 ARATH
ENDO ADPG1 ARATH
ENDO ADPG2 ARATH
ENDO PGLR SOLLC
ENDO QRT2 ARATH
ENDO QRT3 ARATH
C
Arabidopsis thaliana EndoPG protein structure (7B7A)
AT1G05650 model
AT1G05660 model
D
Yersinia enterocolitica ExoPG protein structure (2UVE)
AT1G05650 model
AT1G05660 model

**Figure S5**

A) Evolutionary circle tree showing phylogenetic relationships between plant and non-plant Exo-PGs and Endo-PGs.

B) Evolutionary tree showing phylogenetic relationships between plant Exo-PGs and Endo- PGs with PGRA1 and PGRA2 highlighted by bold text.

C) Protein structure alignment of PGRA1 (yellow) and PGRA2 (light blue) models with plant Endopolygalacturonase structure (orange) resolved for At5g14650 (PDB code 7B7A).

D) Protein structure alignment of PGRA1 (yellow) and PGRA2 (light blue) models with *Yersinia enterocolitica* Exopolygalacturonase structure (pink; PDB code 2UVE). For both (C) and (D) sections, structure (1) corresponds to structure template to which PGRA1 (2) and PGRA (3) were aligned. Alignment (4) shows all three structures in the alignment, whereas (5) and (6) corresponds to the template-model alignment of PGRA1 and PGRA2, respectively. Organism shortcuts used in the phylogenetic trees are ARATH (*Arabidopsis thaliana*), ASPAC (*Aspergillus aculeatus*), ASPAW (*Aspergillus awamori*), ASPFA (*Aspergillus flavus*), ASPFC (*Aspergillus fumigatus*), ASPFN (*Aspergillus flavus*), ASPFU (*Aspergillus fumigatus*), ASPNA (*Aspergillus niger*), ASPNC (*Aspergillus niger*), ASPNG (*Aspergillus niger*), ASPOR (*Aspergillus oryzae*), ASPTN (*Aspergillus terreus*), ASPTU (*Aspergillus tubingensis*), BACSU (*Bacillus subtilis*), COCCA (*Cochliobolus carbonum*), EMENI (*Emericella nidulans*), GIBFU (*Gibberella fujikuroi*), GOSHI (*Gossypium hirsutum*), KALTU (*Kali turgidum*), LACP7 (*Lachnoclostridium phytofermentans*), MAIZE (*Zea mays*), NEOFI (*Neosartorya fischeri*), OENOR (*Oenothera organensis*), PECCC (*Pectobacterium carotovorum*), PECPM (*Pectobacterium parmentieri*), PLAAC (*Platanus acerifolia*), RALSL (*Ralstonia solanacearum*), RHIO9 (*Rhizopus delemar*), SOLLC (*Solanum lycopersicum*), THEIA (*Thermoanaerobacter italicus*), THESW (*Thermoanaerobacterium saccharolyticum*), YEAST (*Saccharomyces cerevisiae*).

A

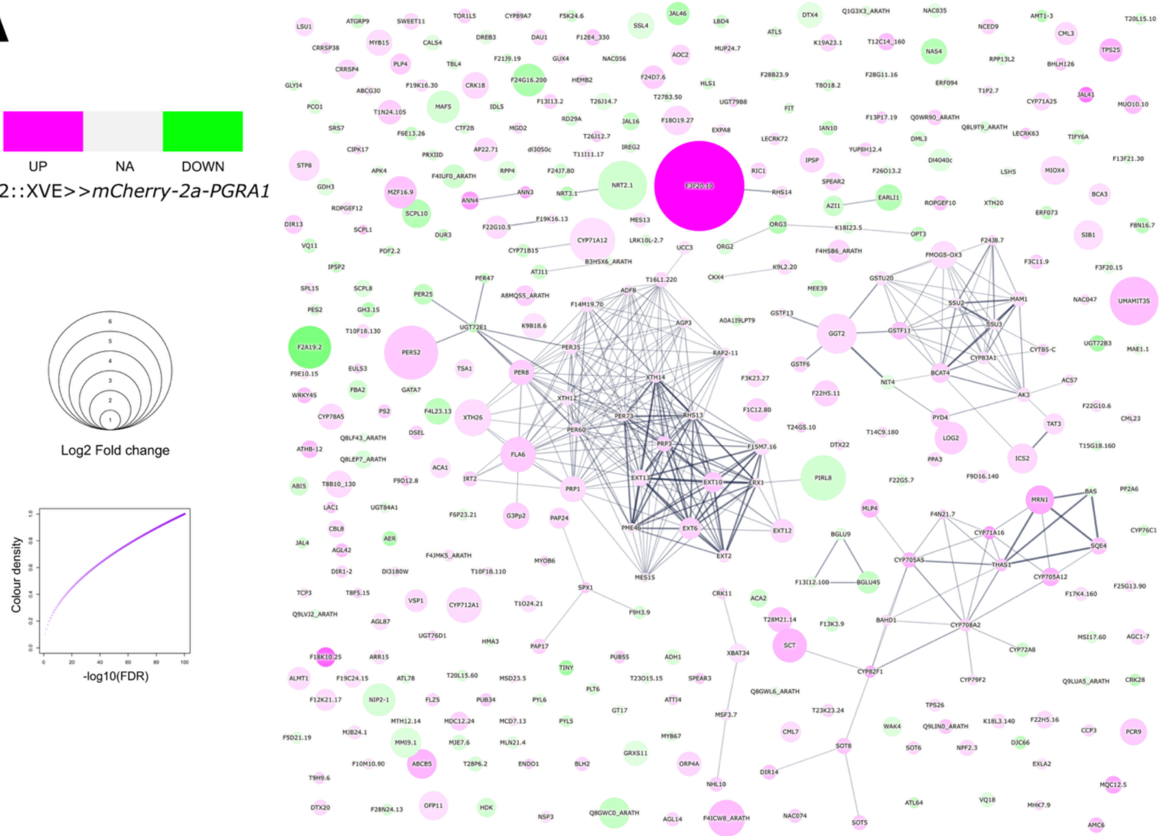
UP
NA
DOWN
PIN2::XVE>>mCherry-2a-PGRA1
Log2 Fold change
Colour density
-log10(FDR)
F3F20.10

B

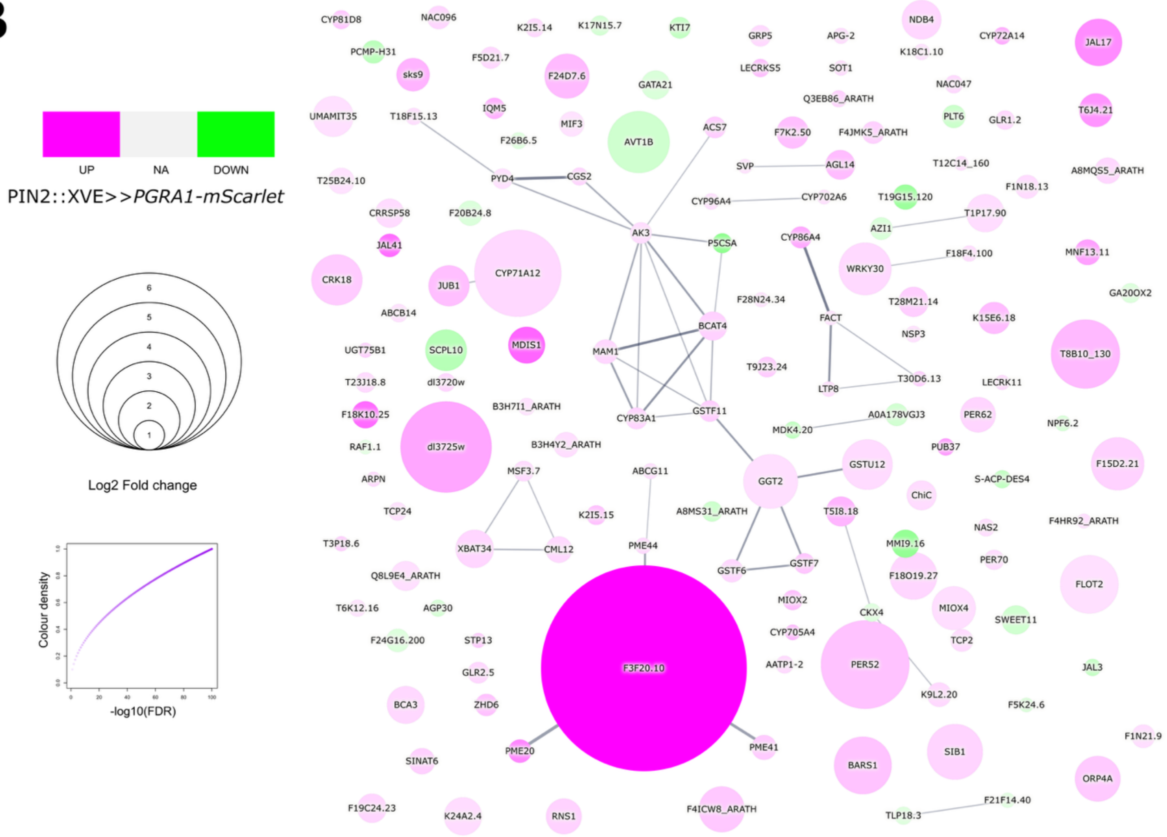
UP
NA
DOWN
PIN2::XVE>>PGRA1-mScarlet
Log2 Fold change
Colour density
-log10(FDR)
CYP81D8
NAC096
K2I5.14
K17N15.7
KTI7
GRP5
APG-2
NDB4
CYP72A14
JAL17
PCMP-H31
F5D21.7
LECRKS5
SOT1
K18C1.10
sks9
F24D7.6
GATA21
NAC047
Q3EB86_ARATH
UMAMIT35
T18F15.13
IQM5
MIF3
PLT6
GLR1.2
T6J4.21
AVT1B
ACS7
F7K2.50
F4JMK5_ARATH
F26B6.5
SVP
AGL14
T12C14_160
A8MQS5_ARATH
T25B24.10
PYD4
CGS2
F1N18.13
CYP96A4
CYP702A6
T19G15.120
CRRSP58
F20B24.8
AK3
T1P17.90
JAL41
P5CSA
AZI1
CYP86A4
MNF13.11
F18F4.100
CRK18
JUB1
CYP71A12
WRKY30
GA20OX2
F28N24.34
ABCB14
T28M21.14
FACT
K15E6.18
BCAT4
NSP3
MDIS1
UGT75B1
SCPL10
MAM1
T8B10_130
T9J23.24
T23J18.8
dl3720w
T30D6.13
LTP8
LECRK11
B3H7I1_ARATH
F18K10.25
A0A178VGJ3
CYP83A1
GSTF11
PER62
MDK4.20
NPF6.2
RAF1.1
dl3725w
B3H4Y2_ARATH
PUB37
ABCG11
GSTU12
F15D2.21
MSF3.7
S-ACP-DES4
ARPN
GGT2
ChiC
K2I5.15
A8MS31_ARATH
T5I8.18
TCP24
F4HR92_ARATH
NAS2
T3P18.6
XBAT34
CML12
PME44
MMI9.16
PER70
GSTF7
GSTF6
Q8L9E4_ARATH
F18O19.27
FLOT2
MIOX2
T6K12.16
AGP30
CKX4
MIOX4
SWEET11
CYP705A4
F24G16.200
STP13
TCP2
F3F20.10
AATP1-2
PER52
JAL3
GLR2.5
K9L2.20
BCA3
ZHD6
F5K24.6
F1N21.9
PME41
SIB1
SINAT6
PME20
BARS1
ORP4A
F21F14.40
F19C24.23
K24A2.4
RNS1
F4ICW8_ARATH
TLP18.3

**Figure S6**

A) STRING network of differentially expressed genes in pPIN2::XVE>>*mCherry-2a-PGRA1* line compared to Col-0 (*FDR* ≤ 0.05 and |*fold change*| ≥ 2). Color coding indicates significant up- or downregulation, circle diameter and color intensity represent *log2FC* and √-log10(*FDR*), respectively. Connection lines indicate the confidence of all interaction types except text-mining. Complete set of DEGs is shown.

B) STRING network of differentially expressed genes in pPIN2::XVE>>*PGRA1-mScarlet* line compared to Col-0 (*FDR* ≤ 0.05 and |*fold change*| ≥ 2). Color coding indicates significant up- or downregulation, circle diameter and color intensity represent *log2FC* and √-log10(*FDR*), respectively. Connection lines indicate the confidence of all interaction types except text-mining. Complete set of DEGs is shown.

**A**

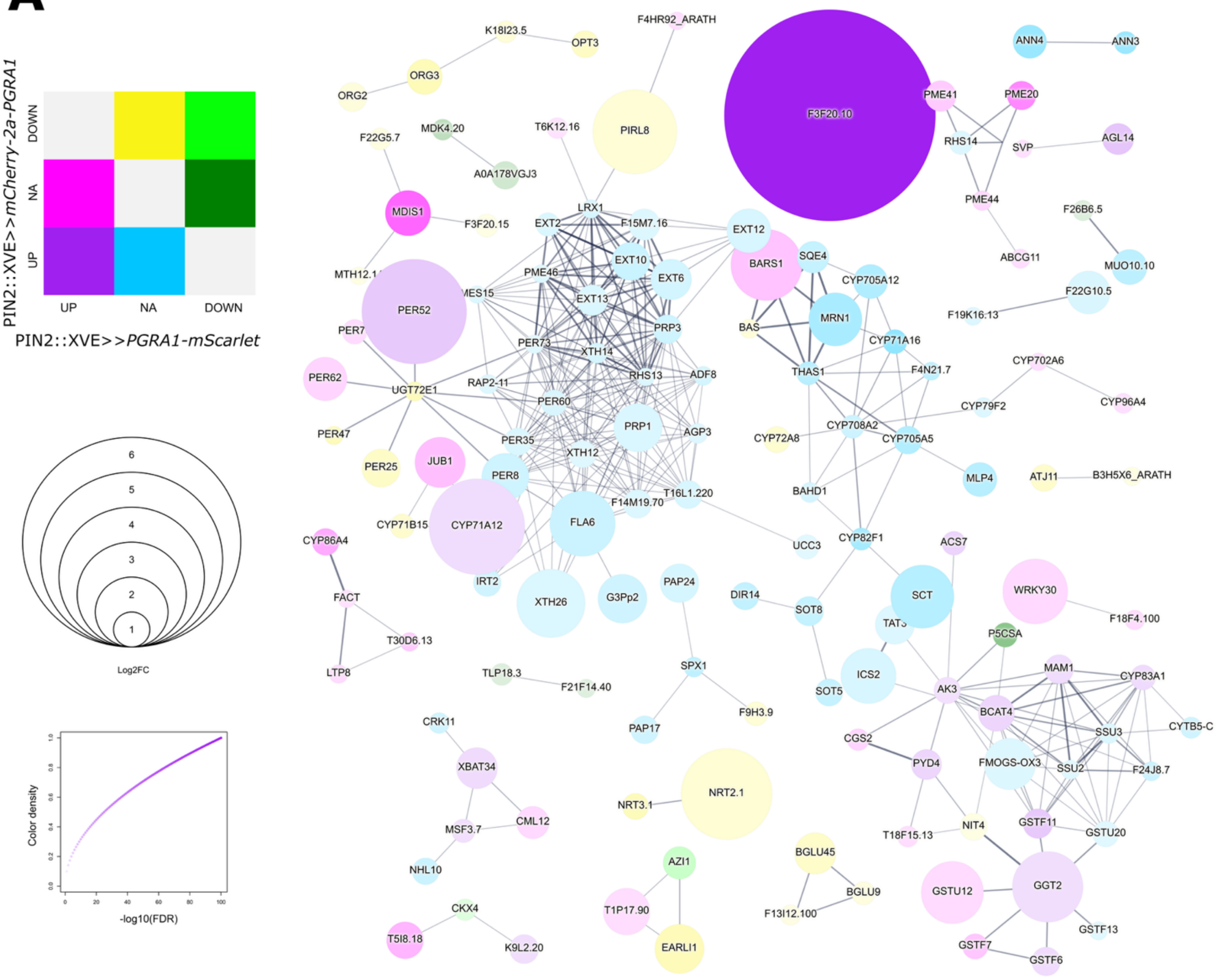


**B**

| GO Term | Fold enrichment | p-value |
|---|---|---|
| GO:0009664~plant-type cell wall organization | 11.61864 | 1.00E-06 |
| GO:0006979~response to oxidative stress | 3.877141 | 7.43E-05 |
| GO:0080003~thalianol metabolic process | 83.91241 | 4.18E-04 |
| GO:0042744~hydrogen peroxide catabolic process | 6.055535 | 0.001063 |
| GO:0009651~response to salt stress | 2.915071 | 0.001134 |
| GO:0009737~response to abscisic acid | 2.724429 | 0.001342 |
| GO:0009625~response to insect | 9.323601 | 0.001976 |
| GO:0009611~response to wounding | 3.525731 | 0.002255 |
| GO:0006952~defense response | 2.302475 | 0.002293 |
| GO:0009407~toxin catabolic process | 8.926852 | 0.002322 |
| GO:0009414~response to water deprivation | 2.818763 | 0.002411 |
| GO:0009413~response to flooding | 35.96246 | 0.002836 |
| GO:0071456~cellular response to hypoxia | 3.343124 | 0.003215 |
| GO:0009409~response to cold | 2.670952 | 0.005668 |
| GO:0019761~glucosinolate biosynthetic process | 10.48905 | 0.006368 |
| GO:0006949~syncytium formation | 17.98123 | 0.011634 |
| GO:0016311~dephosphorylation | 7.991658 | 0.013513 |
| GO:0016104~triterpenoid biosynthetic process | 15.73358 | 0.015105 |
| GO:0001666~response to hypoxia | 7.141482 | 0.01828 |
| GO:0009682~induced systemic resistance | 11.98749 | 0.025426 |
| GO:0010150~leaf senescence | 3.545595 | 0.027439 |
| GO:0015706~nitrate transmembrane transport | 10.9451 | 0.030162 |
| GO:0042542~response to hydrogen peroxide | 5.163841 | 0.042202 |
| GO:0010187~negative regulation of seed germination | 8.680594 | 0.046215 |
| GO:0060586~multicellular iron ion homeostasis | 8.680594 | 0.046215 |
| GO:0019310~inositol catabolic process | 41.9562 | 0.046659 |
| GO:0045944~positive regulation of transcription by RNA po | 3.680369 | 0.047147 |

**Figure S7**

A) STRING network of differentially expressed genes in pPIN2::XVE>>*PGRA1-mScarlet* or pPIN2::XVE>>*mCherry-2a-PGRA1* line compared to Col-0 (*FDR* ≤ 0.05 and |*fold change*| ≥ 2 in at least one comparison). Color coding indicates significant up- or downregulation in one or both lines. Circle diameter and color intensity represent *log2FC* and √-log10(*FDR*), respectively, with the higher values for *FDR* and mean values for *log2FC* among both lines indicated. Connection lines indicate the confidence of all interaction types except text-mining. Non-connected genes are omitted from the interaction network.

B) Gene ontology (GO) terms of differentially expressed genes in pPIN2::XVE>>*PGRA1-mScarlet* or pPIN2::XVE>>*mCherry-2a-PGRA1* line compared to Col-0, grown for 5d on estradiol. p-value ≤ 0.05 and Fold change ≥ 2.

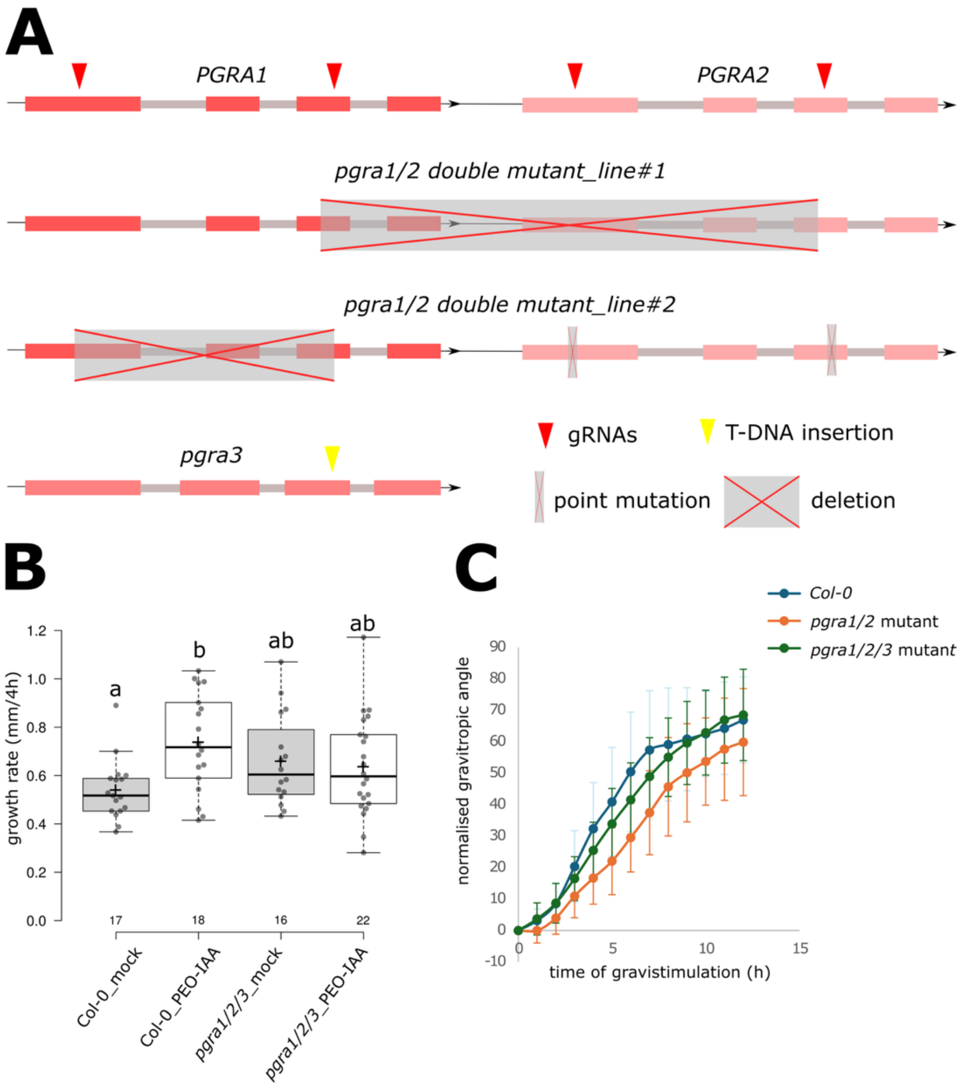


**Figure S8**

A) Schematic representation of the *pgras* mutant lines. The first row illustrates the recognition sites for gRNAs targeting the PGRA1-2 locus. Two versions of *pgra*1/2 mutants are depicted: Version 1 shows a deletion (indicated by a gray rectangle with a red cross) between the end of the *PGRA1* gene and the beginning of the *PGRA2* gene. Version 2 features a deletion within the *PGRA1* gene, along with two point mutations in the *PGRA2* gene. Additionally, the *pgra*3 mutant is shown, with the location of the T-DNA insertion indicated. Exons are in pinkish color, grey color indicate introns.

B) Growth rate of *pgra1/2/3* mutant line treated for 4h with 10µM PEO-IAA.

C) Normalized gravitropic angle of *pgra1/2 and pgra1/2*/3 mutant grown without sugar. n≥ 10.

Different letters indicate statistically significant differences between groups based on one-way ANOVA followed by Tukey HSD (B).

### SUPPLEMENTAL TABLES

#### Table S1: List of gRNAs used for preparation of lines

| target gene | gRNAs Sequence (5´-3´) |
|---|---|
| AT1G05650 | ATATGTACCCTAACACCGGGAGG |
| | ATAATCTTCCGGAGCAACGACGG |
| | AGTGAAATCGATGAACAGCCAGG |
| | GTATTCACCGGATCACAAAACGG |
| | GTTAACAGATTCTCACTGGTCGG |
| AT1G05660 | GTATCGAGTTCGGTATTCACCGG |
| | GTAGAATCGGTGACTCCGTCCGG |

#### Table S2: List of primers used for genotyping

| line | name | Primer Sequence (5´-3´ primers) |
|---|---|---|
| *pgra1/2 double_line 1* | F_AT1G05660 | ATGTCTCGATTAGCGCCTCC |
| | R_AT1G05650 | AACCTTGGTTGGACGGACAA |
| | R_AT1G05660 | AGGTCTTGGAAGAAGACGGT |
| *pgra1/2 double_line 2* | F_AT1G05660 | ATGTCTCGATTAGCGCCTCC |
| | R2_AT1G05660 | CTGATTTGCATCGAATCACCGT |
| GK-343D11 | LB_AT2G43890 | TTTTCGTGGGGGTTTTTAATC |
| | RB_AT2G43890 | AAATACTCTGCGTCCATGACG |
| | LB_GabiKAT | ATATTGACCATCATACTCATTGC |

#### Table S3: List of primers used for cloning

| part | | Primer Sequence (5´-3´ primers) |
|---|---|---|
| PGRA1 promoter | F | GCGCCGTCTCGCTCGGGAGCAGCAGAGAGCTGGCTTTAA |
| | R | GCGCCGTCTCGCTCACATTTTTTTATATGTGAGTTTGAGTTTATGTTT |
| *PGRA1* gene (including introns) B3-B4 position | F | GCGCCGTCTCGCTCGAATGACAAAGTCAGCTATAACATTTCC |
| | R | GCGCCGTCTCGCATCTCCGGTCTGAACGGTG |
| | F | GCGCCGTCTCGGATGATTGTGTTGCTATCGGT |
| | R | GCGCCGTCTCGCTCACGAACCTCGATTTAGACAACTAGTAGGTTG |
| *PGRA1* gene (including introns) B5 position | F | GCGCCGTCTCGCTCGTTCGATGACAAAGTCAGCTATAACATTTCC |
| | R | GCGCCGTCTCGCATCTCCGGTCTGAACGGTG |
| | F | GCGCCGTCTCGGATGATTGTGTTGCTATCGGT |
| | R | GCGCCGTCTCGCTCAAAGCTCATCGATTTAGACAACTAGTAGG |

#### SUPPLEMENTAL METHODS

##### Cell Wall Isolation, Fractionation and ELISA profiling

Protocols for isolation of alcohol insoluble residues (AIR) were previously described (Goubet et al., 2002). Roots of 5d old Col-0, pG1090::XVE>>*ΔMP* and pG1090::XVE>>*AXR3-1* treated for 4h with estradiol were placed in 2 mL tubes with 2 ball bearings, frozen in liquid nitrogen, and ground in the tissue lyser for 2 minutes. Then, 1 mL of 70% ethanol was added to the tube and mixed by rocking for 1 hour at room temperature. After 1 hour, tubes were centrifuged at 10,000 rpm for 15 minutes, and the supernatant was pipetted out. The same procedure was repeated for 80% ethanol, 90% ethanol, and 100% acetone. After the acetone was removed, 1 mL of methanol:chloroform (2:3) was added and incubated for 1 hour. Tubes were then centrifuged at 10,000 rpm for 20 minutes, the supernatant was pipetted out, and the tubes with samples were left open under a fume hood overnight to dry.

Cell wall fractionation for the extracted material was adapted from (Santiago-Doménech et al., 2008). A 50 mM CDTA (pH 6) was used to extract loosely bound homogalacturonans, while 4M KOH was used to extract tightly bound homogalacturonan integrated with other cell wall components (for e.g. xyloglucans). 2 mg of AIR was placed in 2 mL tubes with 2 ball bearings in each tube and ground again with a tissue lyser at 50 Hz for 2 minutes. 1 mL of 50 mM CDTA (pH 6) was added, and the samples were placed in the tissue lyser for 20 minutes, followed by incubation on a rocker at room temperature for 40 minutes. After the incubation, samples were centrifuged at 14,000 rpm for 15 minutes. The supernatant (CDTA extract) was then pipetted out into a new tube and stored at 4°C. For KOH extraction, 1 mL of 4 M KOH and 1% NaBH4 were added to the pellet, and the same procedure was repeated as for CDTA extract.

The ELISA method was adapted from (Pattathil et al., 2010). The 50mM CDTA and 4M KOH extracts obtained from AIR were diluted with 1xPBS (Thermo Fisher Scientific) to 1:10, 1:100, 1:250, 1:500 and 1:1000 (depending on sample) to ensure that the measured signal falls within the linear range of detection. The diluted KOH extract was neutralized to pH 7 using 80% acetic acid before use. 96 well ELISA plates (Thermo Fisher Scientific) were coated with 100 µl of the diluted extracts per well and incubated overnight at 4°C. The following day, plates were washed eight times with tap water, patted dry, and blocked with 200 µl of 5% milk powder (Marvel Original dried skimmed milk) in 1xPBS per well for 1.5 hours at room temperature. After blocking, the plates were washed eight times and patted dry. Primary monoclonal antibodies (Verhertbruggen et al., 2009), were applied at a 1:10 dilution in 3% milk/PBS (100 µl per well) and incubated for 1 hour at room temperature. LM20 was only applied to CDTA extract, since 4M KOH extraction de-esterifies homogalacturonans. Plates were washed again, followed by the addition of secondary antibody (Anti-rat IgG-HRP, Thermo Fisher Scientific) at a 1:1000 dilution in 5% milk/PBS (100 µl per well), and incubated for 1 hour at room temperature. After washing, TMB (3,3′,5,5′-Tetramethylbenzidine) High Sensitivity Substrate solution (BioLegend) was added (100 µl per well), and the reaction was allowed to develop until the blue color appeared (4-8 mins). The reaction was stopped by adding 50 µl of 2.5M sulfuric acid per well, and absorbance was measured at 450 nm using a plate reader (Byonoy Absorbance 96). For each sample, the dilution used for quantification was selected based on being within the linear range — before the signal plateaued and where absorbance still decreased proportionally with dilution.
